# *Chlamydia trachomatis* effector Dre1 interacts with dynactin to reposition host organelles during infection

**DOI:** 10.1101/2022.04.15.488217

**Authors:** Jessica Sherry, Lee Dolat, Eleanor McMahon, Danielle L. Swaney, Robert J. Bastidas, Jeffrey R. Johnson, Raphael H. Valdivia, Nevan J. Krogan, Cherilyn A. Elwell, Joanne N. Engel

## Abstract

*Chlamydia trachomatis* is an obligate intracellular pathogen that replicates within a specialized membrane-bound compartment, called the inclusion. *Chlamydia* species express a unique class of effectors, Incs, which are translocated from the bacteria by a Type III secretion system and are inserted into the inclusion membrane where they modulate the host-bacterium interface. *C. trachomatis* repositions specific host organelles during infection to acquire nutrients and evade host cell surveillance, however the bacterial and host proteins controlling these processes are largely unknown. Here, we identify an interaction between the host dynactin complex and the *C. trachomatis* Inc CT192 (CTL0444), hereafter named Dre1 for <u>D</u>ynactin <u>R</u>ecruiting <u>E</u>ffector 1. We show that dynactin is recruited to the inclusion in a Dre1-dependent manner and that loss of Dre1 diminishes the recruitment of specific host organelles, including the centrosome, mitotic spindle, and Golgi apparatus to the inclusion. Inactivation of Dre1 results in decreased *C. trachomatis* fitness in cell-based assays and in a mouse model of infection. By targeting particular functions of the versatile host dynactin complex, Dre1 facilitates re-arrangement of certain organelles around the growing inclusion. Our work highlights how *C. trachomatis* employs a single effector to evoke specific, large-scale changes in host cell organization that establish an intracellular replicative niche without globally inhibiting host cellular function.

## Introduction

Obligate intracellular pathogens establish a replicative niche within host cells, which often necessitates the rearrangement and repurposing of host cellular structures (Moore and Ouellette, 2014). A subset of intracellular pathogens build and reside within a membrane-bound compartment - essentially constructing a novel organelle within the host cell. And as these pathogens require host-derived nutrients, they must accomplish this task without globally inhibiting host cell trafficking and organelle function, all while evading the host cell’s immune surveillance pathways. Here we elucidate how *Chlamydia trachomatis*, an obligate intracellular pathogen with a dramatically reduced genome, interacts with a single ubiquitous and versatile host protein complex to facilitate the rearrangement of specific host organelles around its replicative niche.

*C. trachomatis* is a Gram-negative bacterial pathogen that is an important cause of disease in humans (Bennett et al., 2020). Specific *C. trachomatis* biovars can infect either the conjunctival epithelium that lines the inside of the eyelid or the mucosal epithelial cells of the urogenital tract. Although infections can be treated with antibiotics, the majority of infections are asymptomatic. No effective vaccine exists, resulting in a high global prevalence of disease. The WHO estimates that over 131 million new *C. trachomatis* infections occur annually (Rey-Ladino et al., 2014; World Health Organization, 2016). Sequelae of untreated infection include blinding trachoma for ocular strains or pelvic inflammatory disease and infertility for urogenital strains (Darville & Hiltke, 2010; Malhotra et al., 2013). *C. trachomatis* infections have been linked with the development of both cervical and ovarian cancer (Smith et al., 2004; Zhu et al., 2016). Uncovering the mechanisms by which *C. trachomatis* establishes its intracellular niche may lead to a better understanding of how this pathogen causes disease and inform the development of targeted therapeutics.

All *Chlamydia* species undergo a biphasic developmental cycle in which the infectious Elementary Body (EB) binds and enters non-phagocytic host cells through receptor mediated endocytosis, and once taken up by the host, transitions to the replicative form, the Reticulate Body (RB) (Bastidas et al., 2013; Chiarelli et al., 2020; Elwell et al., 2016). Following internalization, *Chlamydia* remains within a membrane bound compartment (the inclusion) that it actively modifies to block fusion with the lysosome (Elwell et al., 2016; Moore & Ouellette, 2014). As EBs differentiate to RBs, the growing inclusion traffics along microtubules (MTs) in a dynein-dependent manner to the host centrosome (Grieshaber et al., 2003; Hackstadt et al., 1999). The centrosome is a non-membrane bound organelle attached to the nucleus. It serves as a microtubule organizing center (MTOC) by initiating the assembly of MT networks, and subsequently anchoring and stabilizing these networks (Sanchez & Feldman, 2017). The MTOC, in turn, provides a template that specifies the position of other organelles within the host cell. The inclusion associates tightly with the centrosome throughout the remainder of the *C. trachomatis* replicative cycle. As a result, the centrosome is relocated away from the nucleus during inclusion expansion at late stages of growth (Brown et al., 2014; Grieshaber et al., 2003, 2006; Knowlton et al., 2011). Approximately 24-72 hours following initial infection, RBs differentiate back to EBs, the infectious forms. Mature EBs are released to infect neighboring cells by either host cell lysis or through an exocytosis-like mechanism called extrusion.

*Chlamydia* lacks many essential biosynthetic pathways and must therefore interact with various host compartments such as the cytoskeleton, Golgi Apparatus (GA), mitochondria, lipid droplets, and endoplasmic reticulum (ER) to acquire host-produced metabolites including ATP, sphingolipids, and cholesterol (Dickinson et al., 2019; Elwell et al., 2016). As many of these host cell structures are arrayed around the centrosome, an intimate association with the centrosome would provide the physical opportunity for the inclusion to interact with host compartments and obtain the nutrients required to facilitate intracellular growth. Indeed, *Chlamydia* selectively repositions actin, MTs, the mitotic spindle, GA, ER, and lipid droplets around the developing inclusion during infection, though a detailed understanding of how bacterial and host proteins facilitate this massive host cell reorganization remains elusive (Andersen et al., 2021; Elwell et al., 2016).

*Chlamydia* spp. employ a needle-like Type Three Secretion System (T3SS) to secrete 50-150 effector proteins into the host cytoplasm (Dehoux et al., 2011; Mueller et al., 2014). A subset of these translocated effectors, known as Incs, are transcribed and inserted into the inclusion membrane (IM) at distinct times throughout the *Chlamydia* life cycle (Lutter et al., 2012; Moore & Ouellette, 2014; Weber et al., 2015). Incs are defined by their conserved membrane topology; with two or more short membrane-spanning domains separated by a short loop region (Bannantine et al., 2000). Once inserted into the IM, Incs extend their N-and C-terminal domains into the host cytoplasm (Rockey et al., 2002). Given that Incs are ideally positioned to mediate interactions with the host, we previously utilized an affinity-purification mass spectrometry (AP-MS) based strategy to systematically identify host binding partners of the Incs (Mirrashidi et al., 2015). This study identified a predicted interaction between the *C. trachomatis* Serovar D Inc, CT192 (homolog of CTL0444; hereafter referred to as Dre1 for Dynactin Recruiting Effector 1), an early expressed Inc of unknown function (Almeida et al., 2012), and the host dynactin complex.

Dynactin is critical for most functions of cytoplasmic dynein-1, the primary eukaryotic minus-end directed MT motor. Dynactin serves to link dynein to specific cargo and to enhance processivity of dynein along MTs, facilitating trafficking of various cargo throughout the cell (Holleran et al., 2001; Johansson et al., 2007; Kardon et al., 2009; King & Schroer, 2000; McKenney et al., 2014; Muresan et al., 2001; Reck-Peterson et al., 2018). Together, dynein and dynactin generate the force required to regulate the shape and positioning of various organelles and cellular structures. In addition, dynactin directly binds MTs and functions to anchor and organize MTs arrayed at the centrosome and GA (Corthésy-Theulaz et al., 1992; Lele et al., 2018; Reck-Peterson et al., 2018; Torisawa & Kimura, 2020). Along with many viruses, *C. trachomatis* utilizes dynein to traffic toward the center of the host cell (Grieshaber et al., 2003; Naghavi & Walsh, 2017). In this work, we show that while Dre1 is not required for trafficking of the inclusion along MTs, it instead recruits dynactin to the inclusion to control the positioning of dynactin-positive organelles including the centrosome, mitotic spindle, and GA during infection. By binding a ubiquitous and multifunctional host protein complex Dre1 specifically restructures the host cell interior to facilitate *C. trachomatis* growth without globally inhibiting other host cellular functions.

## Materials and Methods

### Cell culture and bacterial propagation

HeLa 229, Vero, and A2EN cells were obtained from American Type Culture Collection (ATCC). HeLa cells were cultured and maintained in Eagle’s Minimum Essential Medium (MEM; UCSF Cell Culture Facility) supplemented with 10% (v/v) fetal bovine serum (FBS) from Gemini at 37ºC in 5% CO_2_. HEK293T cells (a generous gift from NJ Krogan, UCSF) and Vero cells were cultured and maintained in Dulbecco’s modified Eagle’s Medium (DMEM, UCSF Cell Culture Facility) supplemented with 10% (v/v) FBS at 37ºC in 5% CO_2_. A2EN cells were cultured and maintained in Keratinocyte Media (Gibco) supplemented with 50 µg/mL Bovine Pituitary Extract (Gibco), 0.5 ng/mL Human Recombinant EGF (Gibco), and 10% (v/v) FBS at 37ºC in 5% CO_2_. Cells were routinely tested for mycoplasma (Molecular Probes, M-7006). *C. trachomatis* serovar L2 (434/Bu) and derivative strains used in these studies are listed in Table S3. *C. trachomatis* was routinely propagated in either HeLa 229 epithelial cell monolayers or Vero cell monolayers as previously described (Elwell et al., 2011). HeLa cells were used for all experiments unless otherwise specified. Stellar™ chemically competent *Escherichia coli* (Takara) were used to produce constructs for ectopic expression in mammalian cells, while *dam*^-^/*dcm*^-^ chemically competent *E. coli* (NEB) were used to produce unmethylated constructs for transformation into *C. trachomatis*.

### Plasmid construction

The Dre1 gene and various deletion derivatives used for ectopic expression in mammalian cells were PCR amplified from genomic *C. trachomatis* L2 (434/Bu) DNA and subcloned into the EcoRI and NotI sites in pcDNA4.0/2xStrepII (Jager et. al. 2011) using the primers indicated (Table S3). Dre1 constructs were verified by forward and reverse sequencing. Superfolder (sf) GFP was amplified from a construct kindly provided by Dr. Ron Vale (HHMI, UCSF) and cloned into each Dre1 truncation strain as a C-terminal fusion. Centrin-2 and Mito-7 tagged with mCherry or mEmerald, respectively, were obtained from the Center for Advanced Light Microscopy (Nikon Imaging Center, UCSF). GFP-hARP1a was obtained from the Dumont lab (UCSF). To express epitope-tagged Dre1 during *C. trachomatis* infection, Dre1 was amplified from genomic *C. trachomatis* L2 (434/Bu) DNA and subcloned into the NotI and SalI sites in the *E. coli*/*Chlamydia* pBOMB4 shuttle vector generously provided by Drs. Ted Hackstadt and Mary Weber (Bauler & Hackstadt, 2014). The p2TK2-mCherry *E. coli*/*Chlamydia* shuttle vector encoding pTet-IncG-FLAG was previously generated in collaboration with the Derré lab (Mirrashidi et al., 2015).

### Generation of *C. trachomatis* strains

Rifampin-resistant *C. trachomatis* L2 (434/Bu) was mutagenized using ethyl methanesulfonate to generate a library of nearly 1000 mutants (Nguyen & Valdivia, 2012). Pooled sequencing identified a mutant strain with a single nucleotide variant (SNV) that introduces a stop codon at amino acid 20 of Dre1 (R20*). This mutant was plaque purified (CTL2-M0463, hereafter referred to as L2Δ*dre1*), and subjected to whole genome sequencing as previously described (Kokes et al., 2015). Its DNA sequence was compared to that of the L2 Rif^R^ parental strain to identify other SNVs in L2Δ*dre1* (Table S2). Importantly, L2Δ*dre1* contains no other nonsense mutations. It was re-sequenced periodically to confirm that stocks retained the R20* mutation. Plasmid DNA was isolated from *dam*^-^/*dcm*^-^ *E. coli* or *C. trachomatis* was transformed into *C. trachomatis* L2 Rif^R^ or *C. trachomatis* L2Δ*dre1* as previously described with slight modifications (C. M. Johnson & Fisher, 2013; Y. Wang et al., 2011). Briefly, 10 µg of plasmid was mixed with 1 × 10^7^ infection forming units (IFUs) of *C. trachomatis* L2 in 1X Transformation Buffer (10 mM Tris pH 7.4 in 50 mM CaCl_2_) in 200 µl and incubated at room temperature for 30 minutes. The entire transformation mix was added to Vero cells seeded in 6-well plates (33.3µl/well). At 12 hours post infection (hpi), 5 mg/mL Ampicillin (Sigma) was added to select for transformed *Chlamydia*. After 3 initial passages, Ampicillin was increased to 50 mg/mL until transformed *Chlamydia* was expanded. Clonal populations of transformants were isolated under selection by plaque assay in Vero cells. *C. trachomatis* L2 Rif^R^ (parental strain) and L2Δ*dre1* were each transformed with empty vector. L2Δ*dre1* was transformed with pBOMB4-Dre1_FLAG_ for complementation. The *C. trachomatis* L2 Rif^R^ parental strain was transformed with pBOMB4-Dre1_FLAG_ to generate an overexpression strain (see Table S2 for a list of constructed *C. trachomatis* strains).

### FLAG immunoprecipitations

To generate the Dre1 infection interactome, 8 × 6-well plates of 80% confluent HeLa cells were infected with either L2 expressing plasmid-encoded Dre1_FLAG_ or empty vector at a multiplicity of infection (MOI) of 5 for 36 hours. For all other FLAG immunoprecipitations, 3 × 6-well plates of 80% confluent HeLa cells were infected with the indicated *C. trachomatis* strains expressing a FLAG-tagged Inc for 36 hours. 10 µM MG132 was added 4 hours prior to lysis, and cells were lysed on the plates for 30 minutes at 4ºC in Lysis Buffer (50 mM Tris-HCl pH 7.5, 150 mM NaCl, 1 mM EDTA, 0.5% NP-40, PhosStop, Roche Complete Protease Inhibitor). Lysates were clarified by centrifugation at 13,000 RPM, 4ºC for 15 minutes. Supernatants were then incubated with 30 µl anti-FLAG magnetic beads (Millipore Sigma) rotating overnight at 4ºC. Beads were washed three times in Wash Buffer (50 mM Tris-HCl pH 7.5, 150 mM NaCl, 1 mM EDTA, 0.05% NP-40) and then once in Final Wash Buffer (50 mM Tris-HCl pH 7.5, 150 mM NaCl, 1 mM EDTA). Samples were eluted in 45 µl of Elution Buffer (100 µg/mL FLAG peptide in Final Wash Buffer; Millipore Sigma) for 25 minutes at room temperature with continuous gentle agitation. All purifications were performed in triplicate and assayed by anti-FLAG immunoblot using enhanced chemiluminescence (Amersham Biosciences) or by silver stain (Pierce). For FLAG immunoprecipitations not analyzed by MS, eluates were analyzed by immunoblot analysis with the following antibodies: anti-FLAG, anti-MOMP, anti-GAPDH, and anti-p27.

### Sample preparation and mass spectrometry

Eluates were digested with trypsin for LC-MS/MS analysis. Samples were denatured and reduced in 2M urea, 2 mM DTT, 10 mM NH_4_HCO_3_ for 30 min at 60° C, then alkylated with 2 mM iodoacetamide at room temperature for 45 minutes. Trypsin (Promega) was added at a 1:100 enzyme: substrate ratio and digested at 37° C overnight. Following digestion, samples were then concentrated using C18 ZipTips (Millipore Sigma) according to manufacturer’s instructions. Desalted samples were evaporated to dryness, and resuspended in 0.1% formic acid for MS analysis. Digested peptide mixtures were analyzed by LC-MS/MS on an Orbitrap Fusion Tribrid mass spectrometer (Thermo Fisher Scientific) equipped with an Easy-nLC 1200 HPLC (Thermo Fisher Scientific). Peptides were directly injected onto an analytical column (360 µm O.D. x 75 µm I.D.) with an integrated emitter (New Objective) that was packed with 25 cm of ReproSil Pur C18 AQ 1.9 µm particles (Dr. Maisch). The HPLC system delivered a gradient from 4% to 30% ACN in 0.1% formic acid over 43 minutes, followed by an increase to 80% ACN over 5 minutes, and lastly a hold at 80% ACN for 20 minutes. Peptides were introduced into the mass spectrometer by electrospray ionization in positive mode (1980V) with a transfer tube at 300°C. MS1 scans were performed with orbitrap detection in profile mode at a resolution of 120K, a scan range of 400-1600 m/z, a maximum in injection time of 100 ms, and AGC target of 200K ions, 1 microscan, an S-Lens RV of 60. Peptides with peptide isotopic distribution patterns (MIPS = on) of charge state 2-7 were selected for data-dependent MS2 fragmentation, with a dynamic exclusion time of 20s, a single selection being allowed, a +/-10 ppm mass tolerance, and a minimum signal of 5K. Peptides selected for MS2 were fragmented by beam-type collisional activation (HCD), with a 1.6 m/z isolation window, a first mass of 100 m/z, a collision energy of 30, detection in the ion trap at a rapid scan rate in centroid mode, a 35 ms maximum injection time, and AGC target of 10K, inject ions for all available parallelizable time was activated.

### Proteomics data analysis

The mass spectrometry proteomics data have been deposited to the ProteomeXchange Consortium via the PRIDE partner repository with the dataset identifier PXD028543 (Perez-Riverol et al., 2019). All raw data was searched against both the Dre1 protein sequence and the canonical isoforms of the UniProt human proteome (downloaded June 21, 2021) using MaxQuant (version 1.6.12.0) (Cox & Mann, 2008). Default search parameters were used to with the exception that match between runs was activated with a matching time window of 0.7 minutes. The default parameters included trypsin specificity, a maximum of two missed cleavages, a 1% false discovery rate at the peptide and protein level, a variable modification of oxidation on methionine, a variable modification of acetylation on the protein N-terminus. Finally, protein-protein interaction scoring of the identified proteins was performed with SAINTexpress, and high confidence protein-protein interactions were defined as those with a false discovery rate (BFDR) of less than 10% percent (Teo et al., 2014). We also included previously published high-confidence PPIs for Dre1 with a BFDR < 20% (Mirrashidi et al., 2015).

### Strep affinity purifications

For Strep affinity purifications, approximately 6 × 10^6^ HEK293T cells were seeded in each of three 10 cm^2^ plates, and were transfected using Avalanche-Omni Transfection Reagent (EZ Biosystems), following manufacturer’s instructions. At 48 hours after transfection cells were detached with 10 mM EDTA/D-PBS, washed with PBS, and lysed with 1 mL of ice-cold Lysis Buffer at 4ºC for 30 minutes while rotating. Lysates were incubated with 30 uL of Strep-Tactin Sepharose beads (IBA) in 1 mL of Final Wash Buffer and incubated overnight, rotating at 4°C. Beads were washed three times in 1 mL of Wash Buffer and once in 1 mL of Final Wash Buffer. Samples were eluted in 45 µl of 10 mM _D_-desthiobiotin (IBA) in Final Wash Buffer for 25 minutes at room temperature with continuous gentle agitation. Eluates were immunoblotted with anti-p150^glued^, anti-Strep, and anti-GAPDH antibodies.

### Antibodies and reagents

Primary antibodies were obtained from the following sources: mouse anti-p150^glued^ (BD Biosciences, 610473), mouse anti-FLAG (Millipore, F3165), rabbit anti-FLAG (Millipore, F7425), mouse anti-GAPDH (Millipore, MAB374), mouse anti-GM130 (BD Biosciences, 610823), mouse anti-Centrin (Millipore, 04-1624), mouse anti-dynein, 74 kDa intermediate chains (Millipore, MAB1618), rabbit anti-p27 (DCTN6, Proteintech, 16947-1-AP), rabbit anti-γ-tubulin (Sigma, T3559), rabbit anti-Arl13b (Proteintech, 17711-1-AP), goat anti-MOMP L2 (Fitzgerald, 20C-CR2104GP), rabbit anti-Strep TagII HRP (Novagen, 71591–3), mouse anti-β-tubulin (Sigma, T4026), mouse anti-E-cadherin (Invitrogen, 13-1700), Rat anti-CRB3 (Abcam, ab180835). Mouse anti-IncA and rabbit anti-IncE antibodies were kindly provided by Dan Rockey (Oregon State University) and Ted Hackstadt (Rocky Mountain Laboratories), respectively. Secondary antibodies for immunofluorescence were derived from donkey and purchased from Life Technologies: anti-goat Alexafluor 647, anti-mouse Alexafluor 647, anti-rabbit Alexafluor 647, anti-mouse Alexafluor 568, anti-rabbit Alexafluor 568, anti-rat Alexafluor 568, anti-goat Alexafluor 488, anti-mouse Alexafluor 488, anti-rabbit Alexafluor 488. Nocodazole was purchased from Sigma (M1404). SiR-tubulin 647 was purchased from Cytoskeleton, Inc. Wheat-germ agglutinin 647 was purchased from Life Technologies (W32466). Heparin sodium salt was purchased from Sigma (H3393). (S)-MG132 was purchased from Cayman Chemicals (10012628) and Type 1 Collagen from ThermoFisher (A1048301).

### Fluorescence imaging

HeLa cells were grown on glass coverslips in 24-well plates and infected with the indicated *C. trachomatis* L2 strains (MOI ∼ 1). Bacteria suspended in MEM supplemented with 10% FBS were centrifuged onto cell monolayers at 3500 RPM for 30 minutes at 4°C. Infected cells were incubated at 37°C in 5% CO_2_ for 1 hour, infection media aspirated, and fresh media added. Cells were then incubated at 37°C in 5% CO_2_ for 24, 36, or 48 hours as indicated in the figure legends. For expression of epitope-tagged constructs, HeLa cells were transfected with the indicated constructs using Effectene (QIAGEN) for 24 hours prior to infection, according to manufacturer’s instructions. Experiments requiring inclusion quantitation were performed at a low MOI (∼0.2) to minimize cells with multiple inclusions. Experiments assaying efficiency of inclusion fusion were performed at a high MOI (∼10) to maximize the number of cells with multiple inclusions. When imaging centrosomes or cytoskeletal elements (including dynactin), cells were fixed in 100% ice-cold methanol for 6 minutes. For imaging the GA or transfected fluorescent fusion proteins, cells were fixed in 4% PFA in PBS for 15 minutes at room temperature and then permeabilized in pre-warmed 1X PBS containing 0.1% Triton X-100 for 5 minutes at room temperature. Cells were blocked in 1X PBS containing 1% BSA or 2% BSA (anti-Centrin) for 1.5 hours, and stained with the indicated primary and fluorophore-conjugated secondary antibodies in blocking buffer for 1 hour each. Centrosomes were detected with anti-Centrin to observe centrioles and anti-γ-Tubulin to observe pericentriolar material. MTs and mitotic spindles were stained with anti-β-tubulin or anti-p150^glued^. Of note, there is currently no Dre1 antibody available that detects endogenous levels by immunofluorescence microscopy. Coverslips were mounted in Vectashield mounting media containing DAPI (Vector Laboratories) to identify bacteria and host cell nuclei. When quantitating number of nuclei or inclusions per cell, cells were stained with wheat-germ agglutinin (WGA) 647 to delineate the plasma membrane. To assay Dre1 localization after Nocodazole treatment, HeLa cells were grown on glass coverslips in 24-well plates, transfected with Dre1-sfGFP truncations for 24 hours and then treated with 100 ng/mL Nocodazole or equivalent concentration DMSO for 3 hours, and then cold-shocked for 30 minutes on ice. Cells were then immediately fixed in 4% PFA in PBS for 15 minutes at room temperature, permeabilized in pre-warmed 1X PBS containing 0.1% Triton X-100 for 5 minutes at room temperature and then stained with the indicated antibodies.

For co-localization of transfected Dre1-sfGFP truncation constructs and either (i) the MTOC (stained with the dye SiR-Tubulin 647; Cytoskeleton, Inc) or (ii) the centrosome (co-transfected with mCherry-Centrin2), HeLa cells were seeded on 24-well glass-bottom plates (MatTek, P24G-1.0-13-F) and transfected using Effectene (QIAGEN). At 24 hours post-transfection, cells were incubated with 100 nM SiR-Tubulin 647 dye in MEM for two hours at 37°C in 5% CO_2._ Cells were then stained with 20 nM PureBlu Hoescht (BioRad, 135-1304) in MEM for 15 minutes at 37°C in 5% CO_2_. Cells were then washed with PBS, and incubated in phenol-free DMEM (UCSF Cell Culture Facility) at 37°C and 5% CO_2_ and imaged with a spinning disc confocal microscope (as described below).

### Microscopy

Single Z slices or 0.3 μm-thick Z-stack images were acquired using Yokogawa CSU-X1 spinning disk confocal mounted on a Nikon Eclipse Ti inverted microscope equipped with an Andora Clara digital camera and CFI APO TIRF 60X or 100X oil or PLAN APO 40x objective. Images were acquired by NIS-Elements software 4.10 (Nikon). For each set of experiments, the exposure time for each filter set for all images was identical. Images were processed with Nikon Elements, or Fiji Software.

### A2EN pseudo-polarization

Glass coverslips were placed in 24-well plates and submerged in 500 µl of Type 1 collagen diluted to 30 µg/mL in 20 mM Acetic Acid in ddH_2_O for 5 minutes at room temperature. Following aspiration, coverslips were washed with Keratinocyte media to remove residual acetic acid. Approximately 3.5 × 10^5^ A2EN cells were seeded per well on collagen-coated coverslips. Cells were grown for 48 hours at 37°C in 5% CO2. The media was aspirated to remove non-adherent cells and fresh media was replaced for 24 hours. Cells were infected with L2 strains by centrifuging bacteria diluted into 200 µl of media per well onto cells at 500 g for 5 minutes at room temperature. Infections were aspirated and cells were refed with warm media and incubated for 24 or 48 hours before fixation in either ice-cold MeOH or 4% PFA (see above for further detail). Polarization of cells was confirmed by immunofluorescence imaging of E-cadherin and Crumbs 3.

### Quantitation of inclusion formation

HeLa cells, infected with the indicated L2 strain for 24 hours, were fixed with ice-cold methanol or 4% PFA and visualized by confocal microscopy with anti-MOMP and fluorescent secondary antibodies. Images were acquired using a 40X objective and Nikon Elements pre-assigned image acquisition mode for 6 × 6 fields, creating a minimum of 30 usable fields per coverslip x 3 technical replicates, for a total of ∼90 fields per condition in which to enumerate inclusions. Data are mean ± SD of 3 independent biological replicates. To quantify production of infectious progeny, infected HeLa or A2EN cells were osmotically lysed in ddH_2_O at 24, 36, or 48 hpi. 5-fold serial dilutions of harvested bacteria were used to infect fresh HeLa monolayers. At 24 hpi, inclusion formation was quantified as above.

### Quantitation of centrosome recruitment, centrosome spread and cilia recruitment during interphase

In images of acquired for HeLa cells infected for 36 hours, the centrosome to nucleus distance was calculated using Fiji to create 3D reconstructions of Z-stacks. A line was drawn from the centrosome (stained with anti-γ-Tubulin) to the nearest nuclear face (stained with DAPI) to calculate the centrosome-nucleus distance. Centrosome spread in infected HeLa cells was calculated at 36 hpi by using Fiji to generate maximum intensity projections of 3D image stacks of non-mitotic cells (defined as having uncoiled DNA and lacking spindles). The spread of centrosomes in these projections was calculated using Fiji to draw a polygon encapsulating all centrosomes and then measuring the area of the convex hull corresponding to the polygon. Centrosome spread was calculated in > 40 cells per condition over three independent biological replicates, the average of the three replicates ± SD is overlaid on the individual measurements of centrosome spread. To determine cilia recruitment to the inclusion, we acquired images of A2EN cells grown in serum-containing media, infected for 24 hours, and stained with antibodies to Arl13b and IncA. The percentage of cilia in infected cells with one tip localized within 1 µm of the inclusion membrane was calculated.

### Quantitation of aberrant spindles and multinucleation

HeLa cells infected for 24 hours with the indicated *C. trachomatis* strains were fixed with 100% ice-cold MeOH and stained with antibodies to p150^glued^ and γ-Tubulin to visualize mitotic spindles. The percent of mitotic cells containing aberrant spindles (defined as spindles with > 2 spindle poles) was calculated. For each condition > 45 mitotic cells across 3 independent biological replicates were counted and the average of the three replicates ± SD is represented. Multinucleation rates in infected cells were calculated at 36hpi by staining cells with WGA 647 to delineate the plasma membrane, and the fraction of infected cells containing > 1 nucleus (stained with DAPI) was determined. For each condition > 300 infected cells across 3 independent biological replicates were counted and the average of the three replicates ± SD is represented.

### Quantitation of Golgi Apparatus (GA) recruitment

To calculate GA recruitment to the inclusion membrane, HeLa cells were infected for 24 hours with the indicated *C. trachomatis* strains, fixed with 4% PFA, and stained with antibodies to GM130 (a *cis*-GA marker) and IncE. Fiji was used to generate maximum intensity projections from Z-stacks captured for each condition. Inclusion membrane signal in these 2D projections was traced to form a polygon, which was subsequently fitted to a circle that approximates the inclusion. A second, concentric circle was drawn with a radius 1 µm longer than the inclusion membrane circle to specify the region within the cell that is within 1 µm of the inclusion. The arc length of inclusion membrane corresponding to regions where GA signal falls between the outer and inner circles was transformed to an angle measurement using the angle tool in Fiji, and this value was divided by 360° (see Fig. 5B). For each condition > 100 infected cells across 3 independent biological replicates were counted and the average of the three replicates ± SD is represented.

### Murine Infection Model

All experiments with mice were approved by the Institutional Animal Care and Use Committee of Duke University. Duke University maintains an animal care and use program that is fully accredited by the Association for the Assessment of Accreditation of Laboratory Animal Care, International (AAALAC). Female C57BL/6J mice (Jackson Laboratory) were treated with 2.5 mg medroxyprogesterone (TEVA Pharmaceuticals) subcutaneously to synchronize their estrous cycles. Seven days later, 20 mice were infected transcervically with 1 × 10^7^ EBs per mouse using an NSET Embryo Transfer device (ParaTechs). Mice were sacrificed at 3-and 5-days post infection, and the upper genital tracts were excised and trimmed of adipose tissue and immediately homogenized in 1 mL PBS (Gibco). DNA was extracted using a DNeasy kit (Qiagen) from 80 µL of homogenate following procedures recommended by the manufacturers.

### RT-qPCR

Quantitative PCR was performed on a StepOne Plus Real Time PCR Systems (Applied Biosystems) using Power UP SYBR Green (ThermoFisher Scientific). Quantification of L2 16S rRNA and mouse GAPDH were performed in triplicate and based on standard curves from dilutions of purified *C. trachomatis* and mouse DNA. Mouse PCR targets and primers used were: GAPDH (5’-ACTGAGCAAGAGAGGCCCTA-3’, 5’-TATGGGGGTCTGGGATGGAA-3’), and *C. trachomatis* PCR targets and primers used were: 16S rRNA (5’-GGAGGCTGCAGTCGAGAATCT-3’, 5’-TTACAACCCTAGAGCCTTCATCACA-3’) (Sixt et al., 2017)

### Statistical analysis

For each experiment, 3 or more independent biological replicates were performed and the results are plotted individually or combined and represented as mean ± SD, as described in figure legends. For experiments with naturally high variability due to the asynchronous nature of *C. trachomatis* infections, Superplots were used to represent the data (Figures 3B, 7A, 7B, and 7C; Lord et al., 2020). Briefly, all individual data points from all replicates are represented as small circles and color coded according to replicate number. Average values of each replicate were also color coded, and are represented with triangles. The black horizontal bars represent the average of all replicates ± SD. All statistical analyses were performed using GraphPad Prism 9.0. All assays were analyzed using a one-way ANOVA with a two-tailed Welch’s t-test. *Chlamydia* growth in the murine infection model used a two-tailed Mann–Whitney U-test.

## Results

### Dynactin interacts with Dre1 during *C. trachomatis* infection

Our previous affinity purification-mass spectrometry (AP-MS) screen, referred to here as the “Transfection Interactome”, predicted a high-confidence interaction between Dre1 transiently expressed in HEK293T cells (Figure 1A, B) and all 11 subunits of the mega-dalton sized host dynactin complex (Mirrashidi et al., 2015). Interactions were scored using MiST (Jager et al., 2012) and CompPASS (Sowa et al., 2009) algorithms. These algorithms prioritize interactions based on their reproducibility, abundance, and specificity. Three dynactin subunits that co-purified with Dre1 (CapZa, CapZb, and beta-actin) were not in the top 5% of MiST scores, likely because their specificity scores were penalized due to their interactions with actin filaments which are also regulated by other Incs (Andersen et. al., 2021, Elwell et. al., 2016). They are, however, expected to be biologically relevant binding partners of Dre1.

**Figure 1.**
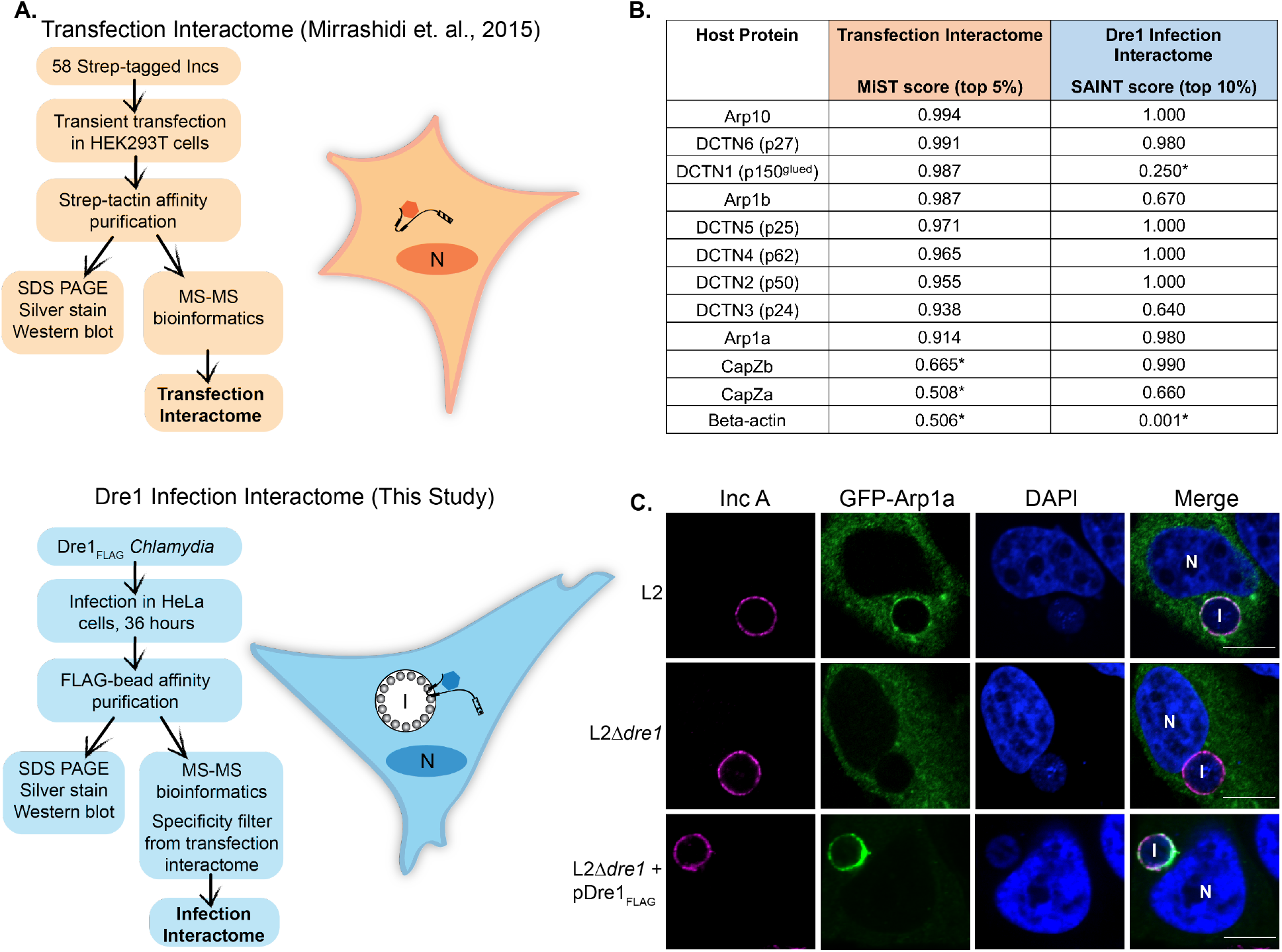
Dre1 recruits host dynactin to the inclusion during infection. (A) Schematic of orthogonal AP-MS screens (“Transfection Interactome” and “Dre1 Infection Interactome”) to identify host binding partners of Dre1. (B) List of dynactin subunits that co-purified with transfected Dre1 in HEK293T cells and scored in the top 5% of all MiST scores (descending order; Transfection Interactome) and of dynactin subunits that co-purified with Dre1 during *C. trachomatis* infection and scored in the top 10% of SAINT scores (Infection Interactome). Host protein scores marked with an asterisk are outside the top 5% or 10% of scores by MiST or SAINT respectively, but are indicated because they were present in Dre1 eluates. (C) Dre1 is required for recruitment of transfected GFP-Arp1a to the inclusion. HeLa cells transfected with GFP-Arp1a (a dynactin subunit) and infected for 24 hours with the indicated strains were fixed and stained with anti-IncA (outlines inclusion membrane) and counter-stained with DAPI (to visualize nucleus and bacteria). Shown are single Z slices. N, nucleus. I, Inclusion. Scale bar, 10µm.

Our previously published Transfection Interactome was performed with Inc proteins from *C. trachomatis* Serovar D, which are highly conserved in Serovar L2 (Dehoux et al., 2011; Lutter et al., 2012). Since *Chlamydia* genetics has been primarily developed for L2, we performed all subsequent studies with L2. To confirm that dynactin interacts with Dre1 in the context of L2 infection and to potentially uncover additional Dre1-interacting partners that may have been missed in the Transfection Interactome, we infected HeLa cells for 40 hours with L2 transformed with a plasmid constitutively expressing Dre1 fused to a FLAG tag (L2+pDre1_FLAG_) and performed affinity purification using anti-FLAG magnetic beads (Figure 1A). Cells infected with L2+pBOMB_FLAG_ vector served as a control. Entire eluates were then subjected to MS analysis. All APs were performed in triplicate, and expression of Dre1 in the eluates was confirmed by immunoblotting with an anti-FLAG antibody and by silver stain (Figure S1A). To generate a set of high-confidence Dre1 interacting partners, we measured the enrichment of host peptides that eluted with Dre1_FLAG_ expressed in L2 during infection compared to control eluates and then scored the reproducibility and abundance of these peptides using the SAINT algorithm (Table 1; Teo et. al., 2014). Next, we compared this data set to the high-confidence interactions predicted for Dre1 in the Transfection Interactome and selected Dre1-host protein-protein interactions in common to both data sets. This strategy generated a list of high confidence interactors specific to Dre1 that occurred infection (“Dre1 Infection Interactome”; Figure 1A and 1B).

As observed in the Transfection Interactome, all dynactin subunits except beta-actin and p150^glued^ were represented within the top 10% of scored interactions for the Dre1 Infection Interactome (Figure 1B). This finding confirms that dynactin subunits are among the most reproducible and abundant interacting partners of Dre1 during infection. p150^glued^, a dynactin subunit that scored highly in the Transfection Interactome, did not score as highly in the Dre1 Infection Interactome due to post-lysis cleavage, likely by the *Chlamydia* protease CPAF (Tan & Sütterlin, 2014). Notably, in both the Transfection and Infection interactomes, Dre1 did not co-purify with dynein or any of the known adaptors that regulate dynein or dynactin activity.

We further confirmed the specificity of the interaction between Dre1 and dynactin during infection by infecting HeLa cells with L2 strains expressing either plasmid-encoded Dre1_FLAG_ or IncG_FLAG_ (an unrelated Inc that does not bind dynactin) and performing FLAG APs. Endogenous dynactin (p27) co-purified with Dre1 but not with IncG (Figure S1B). Together, these data demonstrate that dynactin interacts reproducibly and specifically with Dre1 and that this interaction occurs during *C. trachomatis* infection.

### Dre1 is required for the recruitment of dynactin to the inclusion during infection

Dynactin localizes at multiple sites in the cell, including MTs, the mitotic spindle, centrosome, nuclear envelope, GA, kinetochores, and the cell cortex (Tirumala & Ananthanarayanan, 2020). In HeLa cells infected with L2+pDre1_FLAG_, endogenous p150^glued^ and Dre1 colocalize at the inclusion membrane (Figure S1D). Likewise, transfected GFP-Arp1a is recruited to the inclusion membrane at 24 hours post infection (hpi; Figure 1C). To determine if Dre1 is required to recruit dynactin to the inclusion, we utilized a chemically mutagenized strain of *C. trachomatis* L2 that contains a single nucleotide variant (SNV) in Dre1 that introduces a stop codon at amino acid 20 (Table 2; hereafter we refer to this mutant as L2*Δdre1*). Even if the 20 amino acid peptide is expressed and is stable, it is not predicted to bind dynactin (see Figure 2). Indeed, in HeLa cells infected with L2Δ*dre1*, GFP-Arp1a is not recruited to the inclusion. To definitively link loss of Dre1 with loss of dynactin recruitment, we complemented L2Δ*dre1* with a plasmid constitutively expressing Dre1 (L2Δ*dre1*+pDre1_FLAG_). Recruitment of transfected GFP-Arp1a was restored in the complemented mutant (Figure 1C). Thus, Dre1 is necessary for recruitment of dynactin to the inclusion. These mutants did not display any obvious defect in trafficking of inclusions to the MTOC (Figure 1C), nor did we see a defect in number of inclusions formed as compared to cells infected with wild type L2 (Figure S6A).

**Figure 2.**
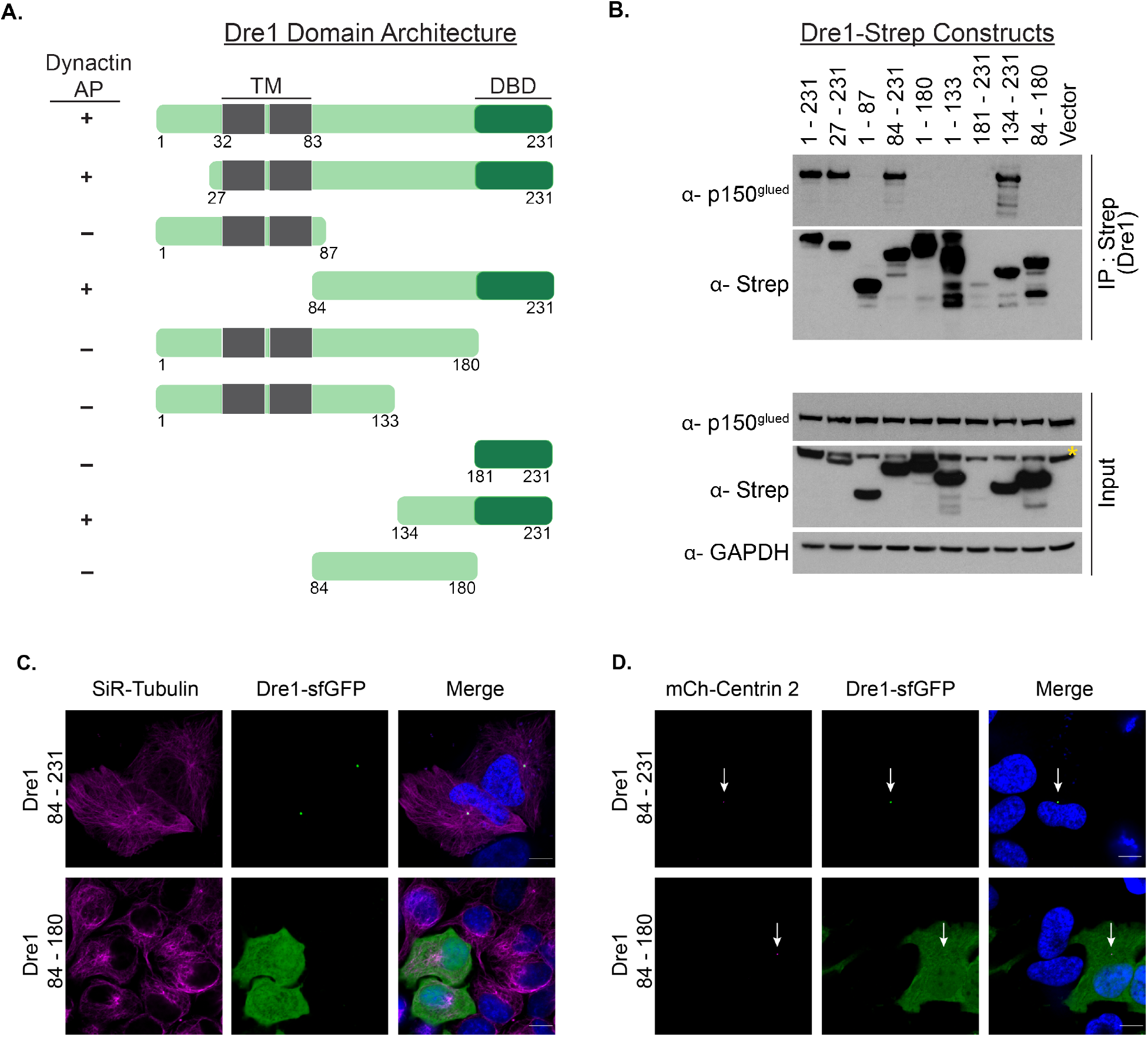
The C-terminal 50 amino acids of Dre1 are required for dynactin interaction and for recruitment of Dre1 to the centrosomal MTOC. (A) Schematic of Dre1_Strep_ constructs and summary of whether they AP with p150^glued^. Transmembrane (TM) domain, grey. Dre1 Dynactin binding domain, (DBD), dark green. (B) Immunoblot of Dre1_Strep_ APs. HEK293T cells were transiently transfected with the indicated Dre1_Strep_ constructs. Lysates were affinity purified with anti-Strep beads, and immunoblotted with the indicated antibodies. Input represents 0.02% of lysates. Cells transfected with empty vector serve as a negative control. Data shown are representative of three independent biological experiments. The asterisk indicates a non-specific band found only in lysates. Since Dre1_Strep_ spanning residues 181-231 did not express well, we cannot definitively assess whether this fragment is sufficient to bind to Dynactin. (C, D) The C-terminus of transfected Dre1 is necessary for co-localization with Tubulin and Centrin 2. HeLa cells were transiently transfected with the indicated Dre1 constructs fused to superfolder GFP (sfGFP), counter-stained with DAPI, and (C) stained with SiR-Tubulin dye to visualize MTs, or (D) co-transfected with mCherry-Centrin 2 to visualize the centrosome (indicated with white arrows). Single Z slices acquired by live cell imaging are shown. Scale bar, 10µm.

### <u>D</u>ynactin <u>B</u>inding <u>D</u>omain targets Dre1 to the centrosomal MTOC

To define the region(s) in Dre1 necessary and sufficient to interact with dynactin, we transfected HEK293T cells with deletion mutants of Dre1. All constructs containing a fragment of the Dre1 C-terminal cytoplasmic domain (Dre1_181-231_) co-AP with endogenous p150^glued^, indicating that this region is required for the interaction with dynactin (labeled <u>D</u>ynactin <u>B</u>inding <u>D</u>omain, DBD, Figure 2 A, B). The construct encompassing the Dre1 C-terminal ∼100 amino acids (Dre1_134-231_) was sufficient to interact with dynactin. Bioinformatic analysis of Dre1_134-231_ failed to reveal any known motifs or any sequence homology to other proteins.

Dynactin localizes to multiple compartments and structures in cells where it organizes MTs, facilitates vesicle trafficking, and positions organelles, suggesting that there may be functionally distinct sub-populations of dynactin (Schroer & Verma, 2021; Tirumala & Ananthanarayanan, 2020). Since *C. trachomatis* inclusions associate with centrosomes and the MTOC (Grieshaber et al., 2003, 2006; Hackstadt et al., 1999), we were particularly interested in the population of dynactin that localizes at the centrosome, where it anchors MTs and organizes MT arrays during interphase and mitosis (Askham et al., 2002; Quintyne et al., 1999; Quintyne & Schroer, 2002). Indeed, live-cell microscopy of HeLa cells showed that the transiently expressed C-terminal fragment of Dre1 (Dre1_84-231_) fused to superfolder GFP (sfGFP) localizes specifically at the MTOC (Figure 2C) and at the centrosome (Figure 2D) but not along MTs. Dre1 localization requires its dynactin-binding domain, as Dre1_84-180_ fails to localize to the MTOC or centrosome (Fig 2C). Furthermore, Dre1 localization does not depend on an intact MT network, as Dre1 and dynactin remain localized at the centrosome in HeLa cells that have been cold-treated with Nocodazole to disrupt microtubules (Figure S2A). Taken together, these data demonstrate that Dre1 specifically interacts with dynactin at centrosomes. We posit that the Dre1:dynactin interaction targets dynactin sub-populations that are involved in organizing MTs nucleated by organelles rather than sub-populations actively involved in MT transport.

### Dre1 repositions centrosomes and primary cilia during infection

In addition to its role in MT organization at the centrosome, dynactin is also involved in centrosome positioning at the juxta-nuclear region within a cell (Burakov et al., 2003; Quintyne et al., 1999). During *C. trachomatis* infection, centrosomes are recruited away from their canonical juxta-nuclear position and instead, maintain a tight association with the inclusion membrane (Brown et al., 2014). This process is dependent on de novo bacterial protein synthesis, suggesting that a *C. trachomatis* effector(s) is involved (Grieshaber et al., 2003). We therefore tested whether Dre1 is necessary for *C. trachomatis*-mediated recruitment of centrosomes to the inclusion.

We performed immunofluorescence (IF) microscopy in HeLa cells infected with L2, L2Δ*dre1*, or L2Δ*dre1*+pDre1_FLAG_ for 36 hours and measured the distance between the centrosomes and the nearest nuclear face in three dimensions (Figure 3A, B). Centrosomes were visualized using antibodies to endogenous Centrin, which stains centrioles. In uninfected cells, the average distance between the centrosome and the nucleus was 0.76 µm. That distance increased to 7.34 µm or 8.66 µm in cells infected with L2 or L2Δ*dre1*+pDre1_FLAG_, respectively. Importantly, in HeLa cells infected with L2Δ*dre1*, the distance between the centrosome and the nucleus was significantly shorter (2.99 µm). Thus, Dre1 contributes to centrosome repositioning during infection, though other *C. trachomatis* effectors may be involved.

**Figure 3.**
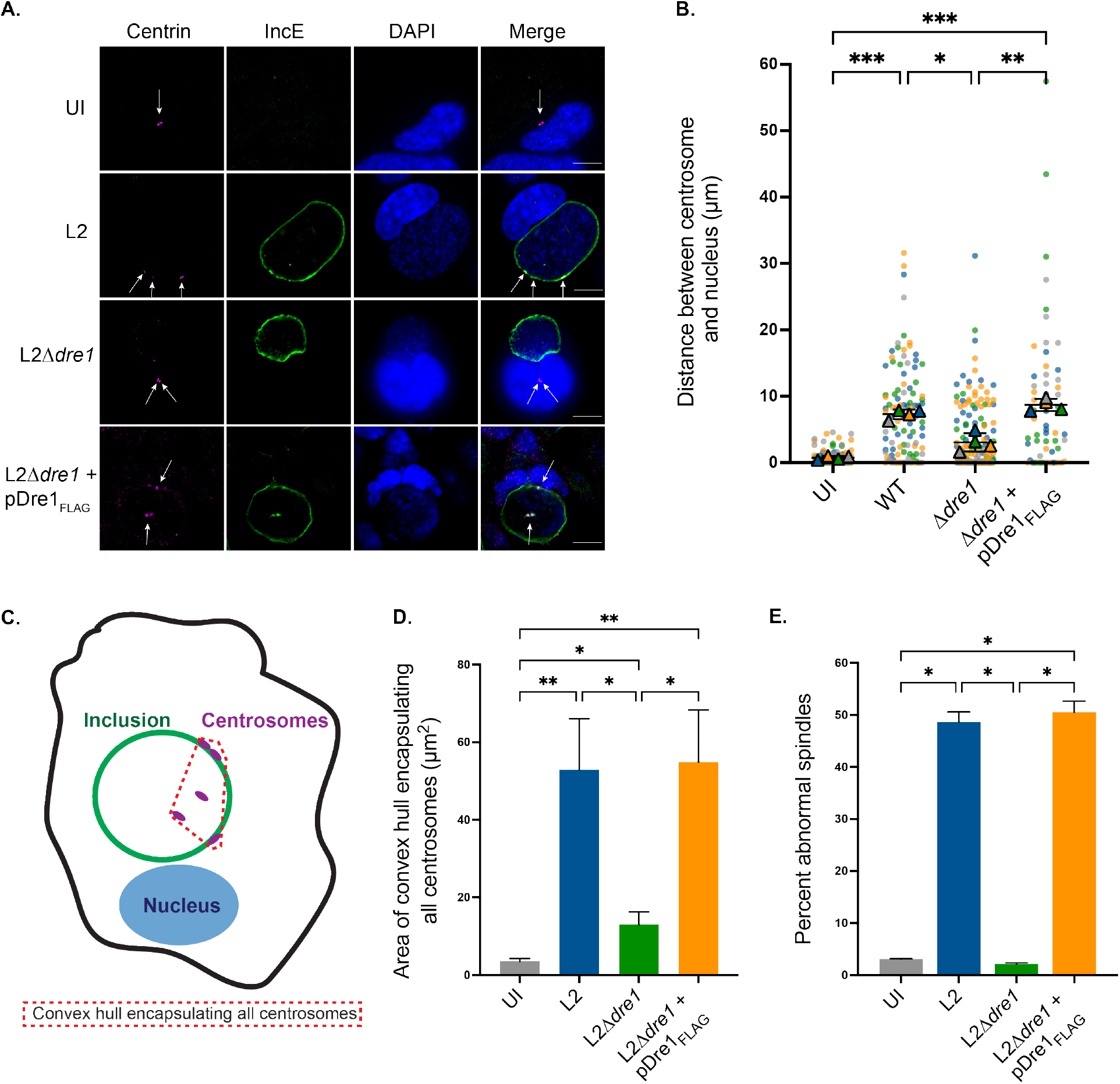
*C. trachomatis* modulates centrosome positioning during infection through Dre1. (A) HeLa cells were infected for 36 hours with the indicated strains or left uninfected (UI), fixed and co-stained with antibodies specific to Centrin (centrosome marker; magenta) and IncE (inclusion membrane marker, green), and counter-stained with DAPI. Shown are single Z slices, arrows indicate centrosome position. Scale bar, 10 µm. (B) Centrosome to nucleus distance in L2-infected cells at 36 hpi was calculated from 3D reconstructions of Z-stacks. Data are represented as individual values for centrosome:nucleus distance for each strain (small colored circles) and average distance for each of three independent biological replicates are represented as triangles (color coded by replicate). Overall average ± SD amongst biological replicates are represented as black bars. 300 cells per condition were counted over three biological replicates. (C) Schematic depicting measurement of centrosome spread. HeLa cells were infected with the indicated strains for 36 hours. Centrosome spread in interphase HeLa cells with > 1 centrosome was calculated by generating maximum intensity projections of 3D image stacks of non-mitotic cells, drawing a polygon connecting all centrosomes and then measuring the area of the convex hull generated from that polygon. (D) Quantitation of centrosome spread. HeLa cells infected with the indicated strains for 36 hours were stained with antibodies to Centrin or γ-Tubulin, and IncE. Centrosome spread was calculated in > 40 cells per condition, over three independent biological replicates. Data are represented as mean ± SD. (E) Quantitation of percentage of mitotic cells with abnormal spindle formation. HeLa cells infected with the indicated strains for 36 hours were stained with antibodies to p150^glued^ and γ-Tubulin to visualize spindles. Spindles with > 2 poles were scored as abnormal. 50 mitotic spindles were analyzed per condition, over three independent biological replicates. For infected samples, only cells with inclusions were quantified. Data are represented as mean ± SD. *p<0.05, **p<0.01, ***p<0.001, Welch’s ANOVA.

The primary cilium is a MT-based sensory organelle involved in the regulation of many cellular processes that originates from the centrosome (Chen et al., 2021). As Dre1 mediates the position of the centrosome during infection, we tested whether Dre1 is involved in positioning of cilia during *C. trachomatis* infection. For these experiments, we utilized A2EN cells, an immortalized human endocervical epithelial cell line that can be grown as a pseudo-polarized monolayer. Pseudo-polarized A2EN cells were grown in the presence of serum, infected, and then fixed and stained for Arl13b (which is highly enriched on the ciliary membrane) and IncA (to delineate the inclusion membrane) at 24 hpi. In L2-or L2Δ*dre1*+pDre1_FLAG_-infected A2EN cells, one end of the cilium localizes at the inclusion membrane in ∼80% of infected cells, while in cells infected with L2Δ*dre1* the base of the cilium is only anchored at the inclusion membrane in ∼15% of cells (Figures S3B, S3C). In cells infected with L2Δ*dre1* the cilium is anchored at the nucleus, which recapitulates the localization of the cilium in uninfected cells. Together these results demonstrate that Dre1 repositions the centrosome as well as the primary cilium, a structure templated by the centrosome, at the inclusion membrane.

### *C. trachomatis* modulates centrosome and spindle positioning during infection through Dre1

In interphase, the centrosome is a single-copy organelle that serves as the primary MTOC in eukaryotic cells. The centrosome is duplicated concurrently with host DNA during S-phase. During M-phase duplicated centrosomes separate from one another and form the two spindle poles that organize MT structures and mediate equal partitioning of host DNA to daughter cells (Bettencourt-Dias & Glover, 2007; Conduit et al., 2015). Migration of duplicated centrosomes depends on dynein and dynactin function (Robinson et al., 1999). Formation of a bipolar spindle is important to avoid chromosome segregation errors and genomic instability. Many cancer cells have supernumerary centrosomes, and in order to successfully replicate, these cells cluster excess centrosomes during interphase and mitosis to form pseudo-bipolar spindles via interactions between the centrosome and MT minus ends (Milunović-Jevtić et al., 2016; Pannu et al., 2014).

*C. trachomatis* infection induces centrosome overduplication, a process reported to be dependent on both the *Chlamydia* protease, CPAF, and on progression of the host centrosome duplication pathway (Brown et al., 2014; Grieshaber et al., 2006; K. A. Johnson et al., 2009). In addition, *C. trachomatis* prevents clustering of these supernumerary centrosomes, which leads to abnormal spindle formation (Brown et al., 2014). To determine whether Dre1 plays a role in centrosome overduplication, we compared the number of centrosomes in uninfected and infected HeLa cells at 36 hpi. While uninfected cells contained on average 2.08 centrosomes per cell, L2-, L2Δ*dre1-* and L2Δ*dre1*+pDre1_FLAG_ -infected cells all exhibited a similar increase in the number of centrosomes per cell (2.76, 2.64, 2.68, respectively, p < 0.05 compared to uninfected cells but p > 0.05 when all L2 strains were compared to each other; Fig S3A). This data indicates that Dre1 is not required for *C. trachomatis*-mediated centrosome overduplication and is consistent with published results showing that while dynactin plays a role in centrosome homeostasis and function, dysregulation of dynactin does not result in centrosome copy number defects (T.-Y. Chen et al., 2015).

We next determined whether Dre1 prevents clustering of supernumerary centrosomes by measuring centrosome spread and spindle polarity in infected HeLa cells at 36 hpi. Cells were stained with antibodies to Centrin or γ-Tubulin, and the number of centrosomes in interphase cells was calculated. Centrosome clustering in cells containing > 1 centrosome during interphase was measured by determining the area of the convex hull that encompasses all centrosomes in a cell (Figure 3C). In L2-and L2Δ*dre1*+pDre1_FLAG_-infected cells, centrosomes occupy an area of 52.9 µm^2^ and 52.5 µm^2^, respectively. In contrast, in L2Δ*dre1*-infected cells, centrosomes occupy an area of 15.7 µm^2^ (Figure 3D). Thus, the Dre1:dynactin interaction at centrosomes overrides the usual clustering mechanisms and enables *C. trachomatis* to dictate centrosome positioning during interphase

Inhibition of centrosome clustering during interphase leads to the development of abnormal spindles during mitosis, which are associated with mitotic failure and/or genomic instability. In addition, dynactin dysregulation induces formation of multipolar spindles (Drosopoulos et al., 2014). We therefore tested whether the Dre1:dynactin interaction contributes to the formation of aberrant spindles during infection. We used a brief cold-shock to synchronize HeLa cells (Rieder & Cole, 2002) and then infected the synchronized cells with L2, L2Δ*dre1*, and L2Δ*dre1*+pDre1_FLAG_ for 24 hours. Cells were fixed and stained with antibodies to γ-Tubulin and p150^glued^ to visualize spindles by IF. Uninfected mitotic cells nearly always formed bipolar spindles, while 47.4% and 52% of mitotic L2-infected or L2Δ*dre1*+pDre1-infected cells exhibited aberrant spindles, respectively (Figure 3E, 4A). Similar to uninfected cells, aberrant spindles were rarely observed in L2Δ*dre1*-infected mitotic cells. These results are consistent with published work suggesting that there are at least two effector pathways that work together to control number and positioning of centrosomes during infection (Brown et al., 2014). Our data support a model where Dre1 positions centrosomes during interphase and mitosis and blocks centrosome clustering but does not dysregulate centrosome copy number.

### Dre1 is required for inclusion localization at the spindle pole during host cell division

Dynactin is recruited to the MT minus ends that congregate at spindle poles during mitosis (Hueschen et al., 2017). Given our results demonstrating that Dre1 mediates association between the inclusion and centrosomes during interphase (Figure 3A), we next tested whether Dre1 maintains this association at spindle poles during mitosis. The *C. trachomatis* inclusion localizes at the center of the spindle and might account for mitotic failure during infection (Greene, 2003; Sun et al., 2011). Dre1-mediated association with dynactin at the spindle pole might be the first step by which the inclusion positions itself with respect to the host spindle to interfere with mitosis.

We determined whether inclusions localize at spindle poles during infection by infecting cold-synchronized HeLa cells for 24 hours and then staining fixed cells for IncE (to delineate the inclusion membrane), p150^glued^ (to reveal the spindle architecture), and DAPI (to visualize chromosomes; Figure 4A). Indeed, L2 inclusions were localized at the spindle poles. In contrast, L2Δ*dre1* inclusions were displaced from the spindle pole and were often found displaced from the spindle and metaphase plate. Association of the inclusion at the spindle pole is restored in the complemented mutant. Interestingly, Dre1-directed localization of the inclusion at the spindle pole often leads to displacement of host chromosomes from the metaphase plate (Figure 4A).

**Figure 4.**
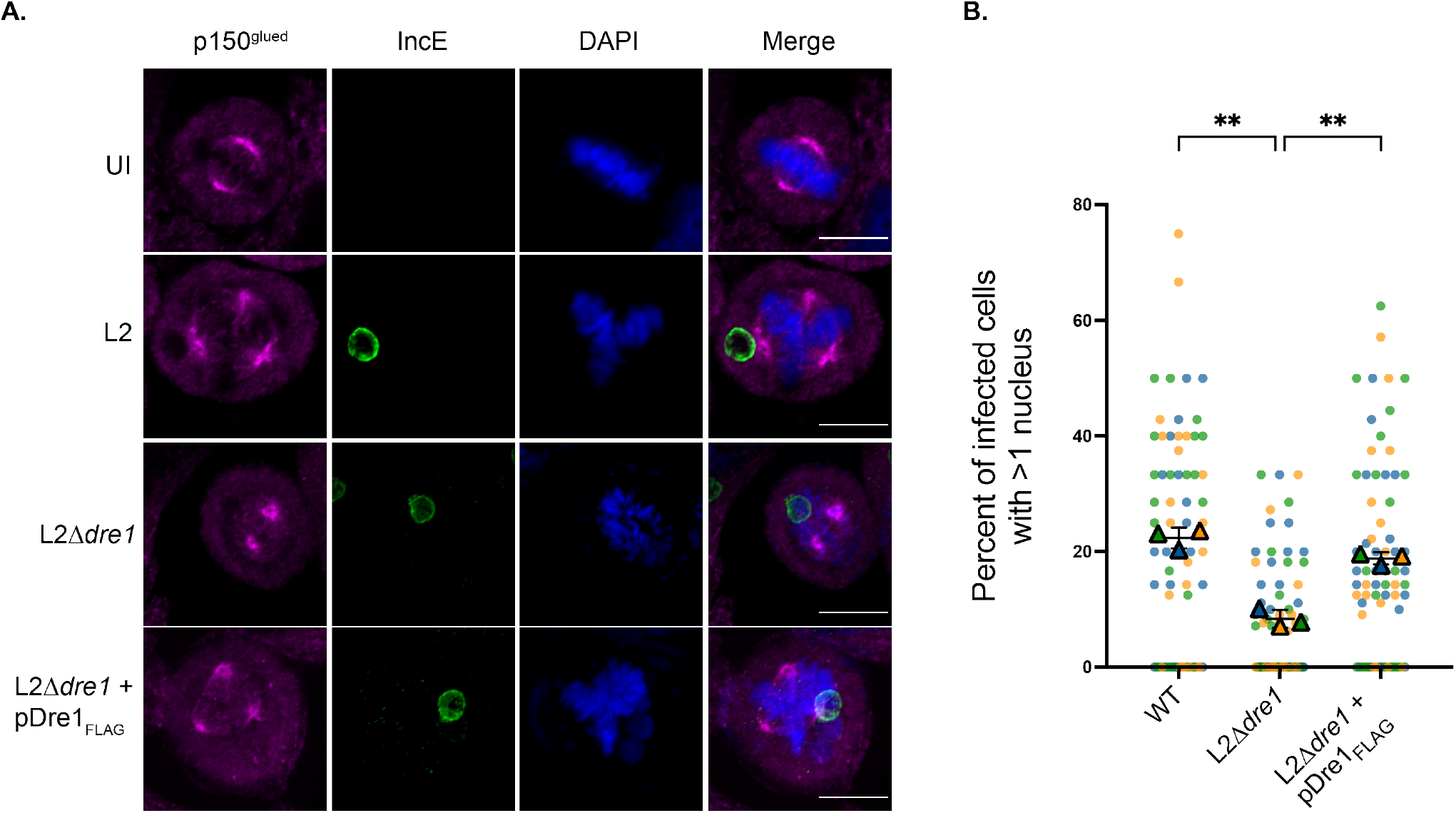
Dre1 positions the inclusion at spindle poles during host division and contributes to *C. trachomatis*-induced multinucleation. (A) HeLa cells were infected with the indicated strains or left uninfected (UI) for 24 hours, fixed and co-stained with antibodies specific to p150^glued^ (to visualize spindle poles, magenta) and IncE (to visualize the inclusion membrane, green), and counter-stained with DAPI. Shown are single Z slices. Scale bar, 10µm. (B) Quantitation of multinucleation. HeLa cells were infected for 48 hours with the indicated L2 strains, fixed and stained with an antibody specific to MOMP (Major Outer Membrane Protein, to visualize bacteria), and counter-stained with DAPI (to visualize nuclei) and WGA 647 (to delineate the plasma membrane). 3D-reconstructions of 75 infected fields were scored for cells containing >1 nucleus for each strain over three replicates. Data are represented as individual percentages for each field (small colored circles) and average percentage for each of three independent biological replicates are represented as triangles (color coded by replicate). Overall average ± SD amongst biological replicates are represented as black bars. **p<0.01, Welch’s ANOVA.

### Dre1 contributes to infection-induced multinucleation

*C. trachomatis* infection induces cytokinesis failure and multinucleation (Alzhanov et al., 2009; Brown et al., 2012; Sun et al., 2016). We therefore tested whether *C. trachomatis* induced host cell multinucleation was Dre1-dependent. Multinucleation was observed in 1.6% of uninfected HeLa cells (Figure S4A), while 22.3% of L2 infected cells were multinucleated by 48 hpi (Figure 4B). In contrast, only 8.3% of cells infected with L2Δ*dre1* were multinucleated, while L2Δ*dre1*+pDre1_FLAG_-infected cells exhibited multinucleation rates similar to L2-infected cells (18.8%). Our results demonstrate that Dre1 plays a role in *C. trachomatis*-induced multinucleation, likely through its effects on centrosome position and spindle architecture.

### Dre1 mediates GA recruitment to the inclusion membrane

Our work demonstrates that Dre1 binds dynactin and modulates the positioning of a MTOC (the centrosome) with respect to the inclusion membrane. In eukaryotic cells, the GA also functions as an MTOC, and dynein/dynactin regulates its structure as well as its perinuclear localization (Jaarsma & Hoogenraad, 2015; Rios, 2014; Yadav & Linstedt, 2011). Disruption of dynactin causes GA fragmentation and dispersal (Palmer et al., 2009; Yadav & Linstedt, 2011). Dynactin can bind to MTs anchored at the GA as well as to βIII spectrin on GA membranes through its Arp1 subunit (Holleran et al., 2001; Yadav & Linstedt, 2011). *C. trachomatis* infection fragments the GA into mini-stacks that are recruited around the inclusion, which enhances progeny production (Heuer et al., 2009). To test the hypothesis that Dre1 mediates recruitment of the GA to the inclusion through its interaction with dynactin, HeLa cells were infected for 24 hours with L2, L2Δ*dre1*, and L2Δ*dre1*+pDre1, fixed, and stained with anti-GM130 (a *cis*-GA marker) (Figure 5A). 52.8% of the inclusion surface area was within 1 µm of the GA in L2-infected cells, whereas only 23.3% of the inclusion surface area in L2Δ*dre1*-infected cells was within 1 µm of the GA (Figure 5C). GA recruitment was restored to L2 levels (55.5%) in the complemented strain. These data demonstrate that Dre1 contributes to GA recruitment at the inclusion membrane.

**Figure 5.**
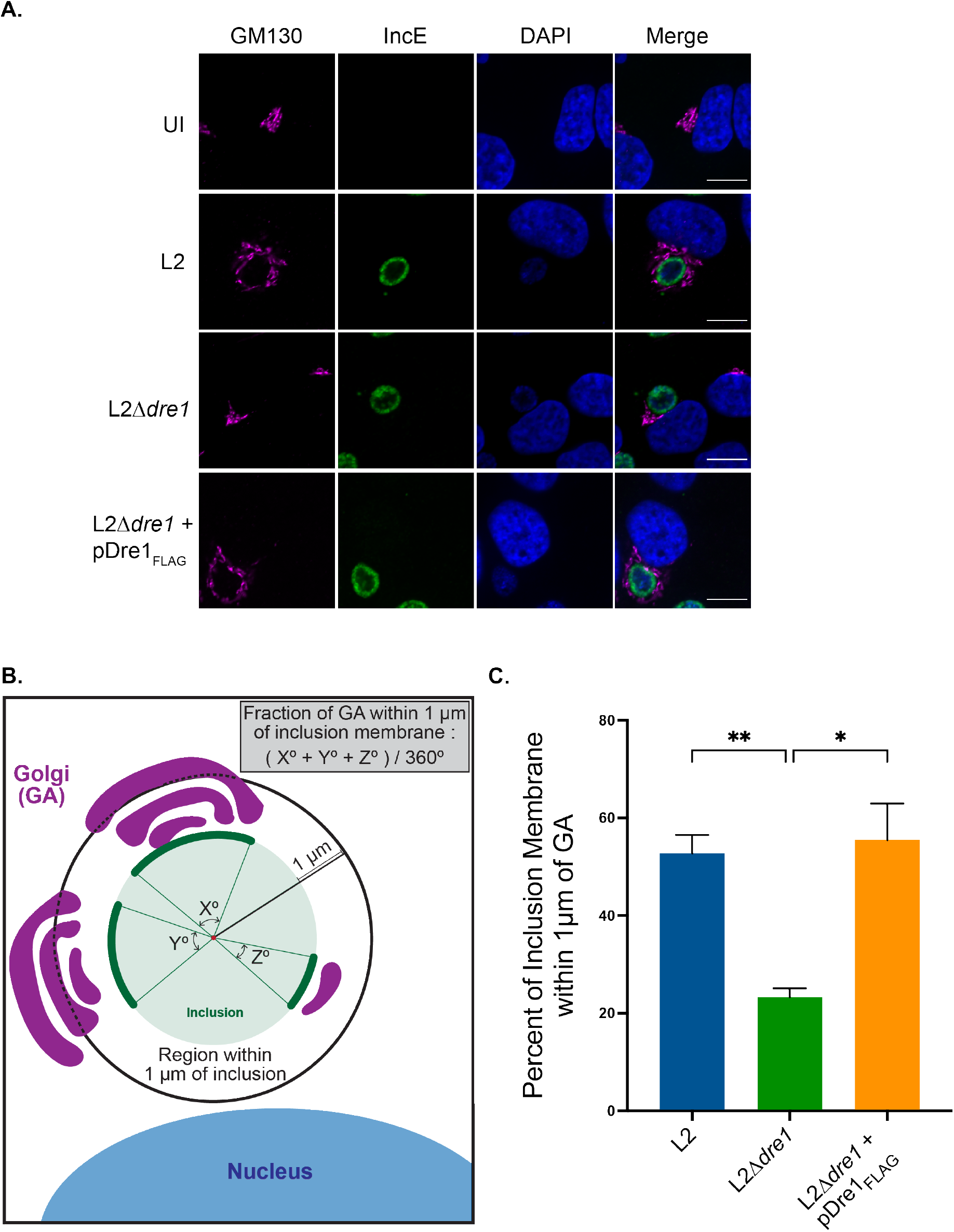
Dre1 is required for the recruitment of the Golgi Apparatus (GA) to the inclusion. (A) HeLa cells were infected with the indicated strains or left uninfected (UI) for 24 hours, fixed and co-stained with antibodies specific to GM130 (to visualize the GA, magenta), IncE (green), and counter-stained with DAPI. Shown are single Z slices. Scale bar, 10µm. (B) Schematic depicting quantitation of GA recruitment to the inclusion. Maximum intensity projections were generated from Z-stacks, and a circle was fitted to the inclusion membrane signal. A second, concentric circle was drawn with a radius 1 µm greater than the inclusion circle. The arc length of inclusion membrane corresponding to regions where GA signal falls between the outer and inner circles was determined, and this value was divided by 360°. (C) Quantitation of GA recruitment in infected cells described in (A). For each condition, > 100 infected cells across 3 independent biological replicates were counted and the average of the three replicates ± SD are represented. *p <0.05, **p<0.01, Welch’s ANOVA.

### Dre1 is required for efficient inclusion fusion at the centrosomal MTOC

In cells infected with multiple L2 bacteria, each bacterium is typically enclosed in a separate inclusion, which then traffics along MTs to the MTOC/juxta-nuclear region. Over a period of ∼ 24 hours, multiple MTOC-localized inclusions undergo homotypic fusion (Richards et al., 2013). While many details of the homotypic fusion process remain to be elucidated, it is known that the inclusion membrane protein IncA is absolutely required for homotypic fusion (Cingolani et al., 2019; Weber et al., 2016). These fusion events contribute to pathogenicity (Geisler et al., 2001) and may be a mechanism for genetic exchange between RBs or a strategy to avoid competing for the same resources in the host cell. Given that inclusion fusion requires association with the centrosomal MTOC (Richards et al., 2013), we tested the hypothesis that Dre1-mediated interaction between multiple inclusions and centrosomal dynactin may facilitate inclusion fusion.

In cells infected with L2Δ*dre1* at a high multiplicity of infection (MOI), we observed a significant delay in inclusion fusion compared to L2 (Figure 6B). At 24 hpi, the average number of inclusions per L2-infected cell was 1.07, indicating that most inclusions had undergone fusion. In contrast, cells infected with L2Δ*dre1* exhibited on average 1.77 inclusions per cell (Figure 6B). The complemented strain had 1.27 inclusions per infected cell, demonstrating that fusion was restored to nearly wild type levels. By 48 hpi, L2Δ*dre1* inclusions are nearly fully fused; thus, Dre1 contributes to efficient inclusion fusion but is not absolutely required. Furthermore, L2-infected cells containing multiple inclusions exhibited Dre1-dependent GFP-Arp1a localization at the boundary membranes between unfused inclusions (Figure 6A). Importantly, loss of Dre1 does not affect IncA expression or localization (Figure 1C).

**Figure 6.**
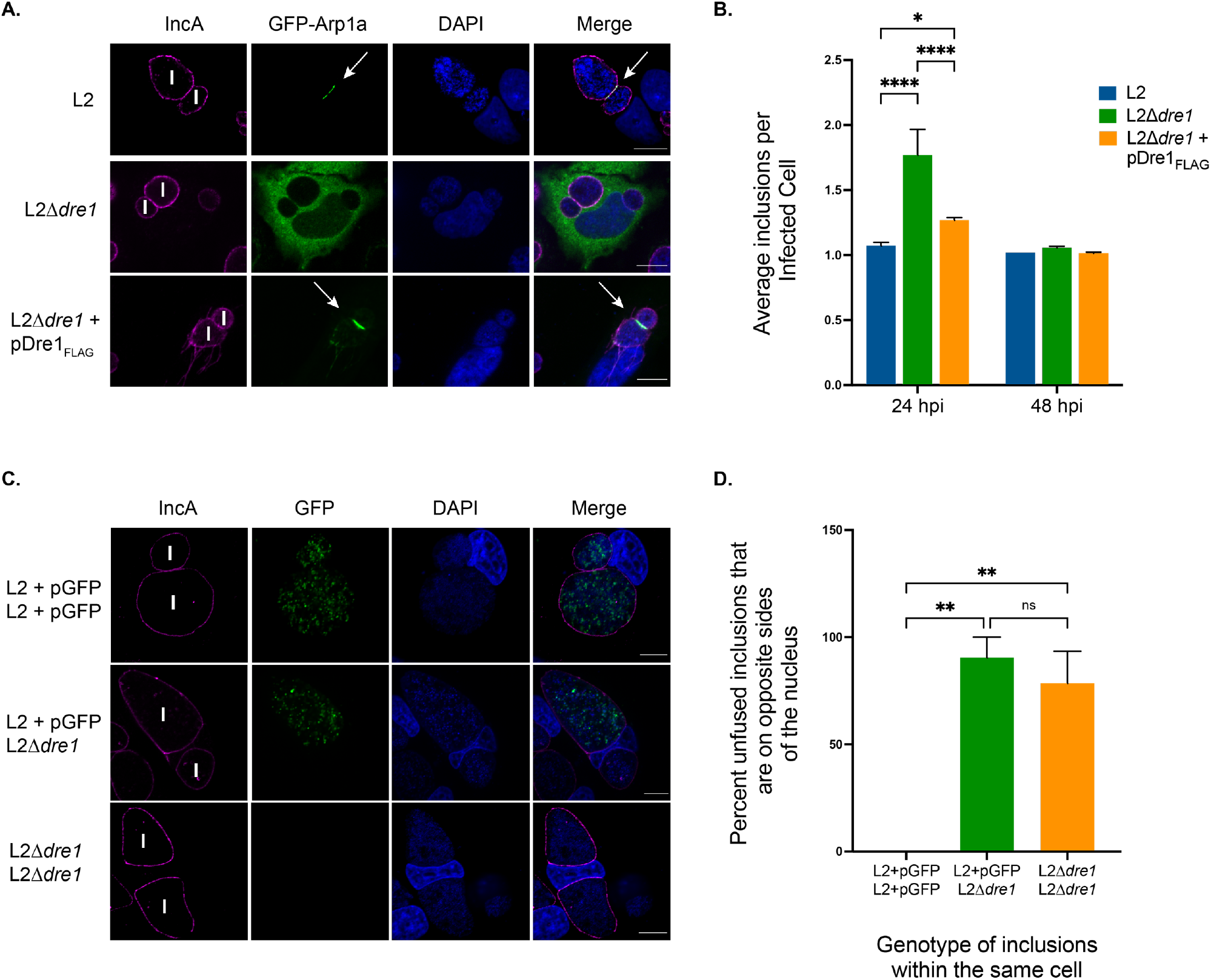
Dre1 is required for efficient inclusion fusion. (A) HeLa cells were transfected with GFP-Arp1a, infected with the indicated strains for 48 hours, fixed, stained with an antibody specific to IncA (magenta), and counter-stained with DAPI. Shown are single Z slices. N, nucleus. I, Inclusion. Scale bar, 10µm. (B) Quantitation of inclusion fusion defect. HeLa cells infected with the indicated strains were fixed at 24 or 48 hpi, stained with an antibody to IncA, and counter-stained for WGA 647 (to outline the plasma membrane), and DAPI. 120 cells were analyzed per condition, over three biological replicates. Data are represented as mean ± SD. (C) HeLa cells were coinfected with L2+pGFP and L2Δ*dre1* for 48 hours fixed, co-stained with an antibody specific to IncA (magenta), and counter-stained with DAPI. Shown are single Z slices. N, nucleus. I, Inclusion. Scale bar, 10µm. (D) Quantitation of infected cells described in (C). 3D-reconstructions of 25 infected fields for each condition over three replicates were scored for cells containing multiple inclusions found on opposite sides of the nucleus from one another. Overall percent ± SD amongst biological replicates are represented as black bars. *p<0.05, **p<0.01, ****p<0.0001, Welch’s ANOVA.

We observed that the rare L2Δ*dre1* inclusions that remained unfused at 48hpi were located on opposite sides of the nucleus from one another (Figure 6A). We hypothesized that these non-fused inclusion events arose as a consequence of trafficking along MTs to minus ends that are not anchored at the primary MTOC. To test this notion, we performed 1:1 coinfections using a GFP-expressing L2 (L2_GFP_), which should localize at the centrosome, and L2Δ*dre1* (which was not fluorescent). At 48 hpi cells were fixed and stained using antibodies to γ-Tubulin and IncA, followed by staining with Wheat Germ Agglutinin (WGA). Figures 6C and 6D show that in the exceedingly rare cases where there are two L2_GFP_ inclusions within the same cell, they are always found immediately adjacent to one another. In a cell containing one L2_GFP_ inclusion and one L2Δ*dre1* inclusion, 90% of inclusions are found on opposite sides of the nucleus. Furthermore, in cells harboring two L2Δ*dre1* inclusions, these inclusions are also found on opposite sides of the nucleus at similar rates as cells harboring both an L2_GFP_ inclusion and an L2Δ*dre1* inclusion. This data supports the idea that the relative defect in L2Δ*dre1* inclusion fusion is at least in part a consequence of the failure of the inclusions to be juxtaposed at the primary MTOC.

### Dre1 is required for virulence in cell culture and a mouse model of upper genital tract infection

Given that L2Δ*dre1* was initially identified by its small plaque phenotype (Kokes et al., 2015), and that loss of Dre1 expression during infection contributes to defects in repositioning host organelles around the inclusion, we tested the contribution of Dre1 to virulence in cell-based and a murine model of upper genital tract infection. First, we quantified whether Dre1 is required for *C. trachomatis* progeny production in HeLa cells or in pseudo-polarized A2EN cells. L2 infections harvested at 48 hpi in HeLa cells, when L2Δ*dre1* no longer exhibits a fusion defect, yielded an 8-fold greater amount of EBs compared L2Δ*dre1* infections (Figure 7B). Likewise, L2 infections at 48 hpi in A2EN cells contain 6-fold more EBs than L2Δ*dre1* (Figure 7A). In both HeLa and A2EN cells, L2Δ*dre1*+pDre1_FLAG_ infection produces comparable numbers of EBs as L2 infection (Figure 7A, B). Thus, Dre1 is required for efficient production of infectious progeny in two different cell lines, and this virulence defect is unlikely due to the delay in inclusion fusion.

**Figure 7.**
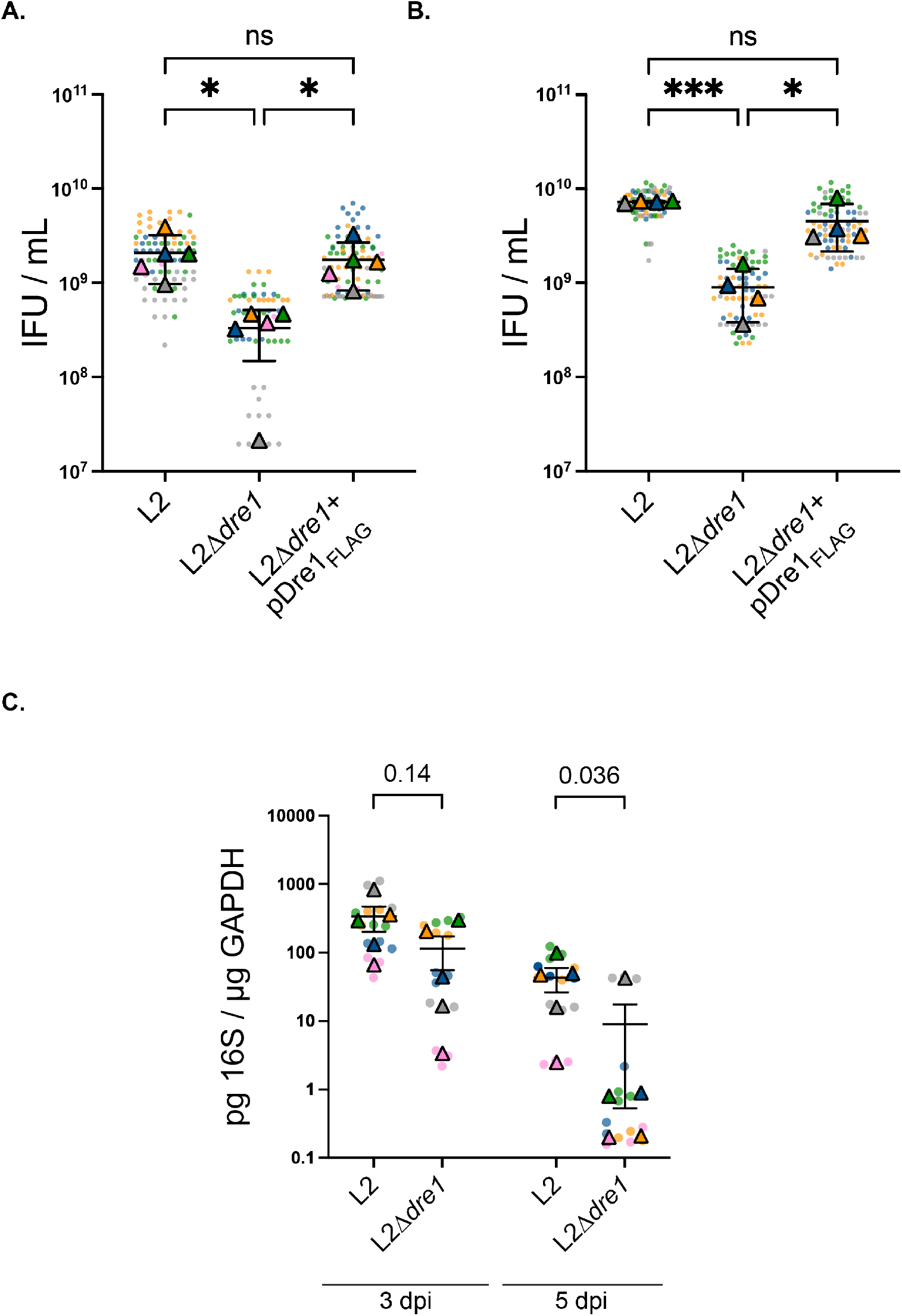
Dre1 is required for virulence in cell culture and a mouse model of upper genital tract infection. (A and B) Quantitation of infectious progeny at 48 hpi in pseudo-polarized A2EN cells (A) or in HeLa cells (B) infected with the indicated L2 strains. Confluent monolayers were infected with the indicated strains for 48 hours. EBs were isolated and used to infect fresh HeLa monolayers to enumerate infectious particles produced over the course of the primary infections. Data are mean ± SD from ≥ 4 independent experiments presented as a scatter plot where all measures (dots), and averages of each biological replicate (triangles) are color coded. *p<0.05, **p<0.01, ***p<0.001, Welch’s ANOVA. (C) Dre1-deficient bacteria are cleared faster from the mouse genital tract, as measured at 3-and 5-days post infection. Data are presented as a scatter plot with technical replicates (dots), and average values for each mouse (triangles) are color coded, n=5, Mann-Whitney U test.

Finally, we tested whether Dre1 contributes to infection in a well-established mouse model of *C. trachomatis*-induced human genital tract disease (Sixt et al., 2017). Female C57BL/6 mice were pre-treated with progesterone for 2 weeks to synchronize their estrus cycles and were then transcervically infected with either L2 or L2Δ*dre1*. This mode of inoculation mimics ascending infection. Mice were sacrificed on days 3 and 5 post infection, and bacterial burden in isolated genital tracts was measured in 5 mice per strain using qRT-PCR against *Chlamydia* 16s rRNA. At 5 days post infection, L2Δ*dre1* was mostly cleared from the mice, while there were 10-fold higher levels of bacteria in the mice infected with L2 (Figure 7C). This result is consistent with the progeny defect observed *in cellulo.*

## Discussion

To evade host-cell innate immune surveillance, internalized *Chlamydia* develop within a membrane-bound compartment. Given that *C. trachomatis* relies on host cell-derived metabolites, this bacterial pathogen avoids globally inhibiting host cell functions while building what is essentially a novel organelle. Through strategic deployment of effectors into the host cytosol and inclusion membrane, *C. trachomatis* remodels host cell structures from within the inclusion. In this work, we identified and characterized the consequences of an interaction between the Inc Dre1 (CT192) and the host dynactin complex (Figure 1). We show that Dre1 interacts selectively with pools of dynactin stably associated with specific organelles including the centrosome, primary cilium, and GA, to modulate their recruitment to the inclusion (Figures 3-5). Importantly, we determined that the Dre1:dynactin interaction is critical to the intracellular survival and pathogenesis of *C. trachomatis* infections (Figure 7). Thus, this single Inc selectively evokes large-scale changes in host cell organelle organization.

Our work underscores the nuances of dynactin regulation and may prove useful in further dissecting its diverse cellular functions. Dynactin is an adaptor for the minus-end directed MT motor dynein, a complex hijacked many intracellular pathogens to facilitate their intracellular transport (Henry et al., 2006). Although *C. trachomatis* is known to traffic along MTs in a dynein-dependent manner to reach the host MTOC, we show that Dre1 is not required for trafficking from the cell periphery to the juxtanuclear position. Furthermore, disruption of dynactin by overexpression of its dynamitin (p50) subunit does not affect trafficking of the inclusion to the centrosome (Grieshaber et al., 2003). These observations suggest that another effector can fulfill this role, a likely candidate being the Inc CT850, which has been shown to interact with dynein (Mital et al., 2015). Given that Dre1 is one of the few pathogen effectors that specifically targets dynactin rather than dynein (Bhavsar et al., 2007; Bouwman et al., 2013; Henry et al., 2006) and that Dre1 does not interact with any known adaptor proteins that enhance dynein processivity (Table 1), we therefore speculate that the primary role of the Dre1:dynactin interaction might be to target the MT-binding and MT-organizing functions of dynactin, rather than targeting actively trafficking dynactin/dynein complexes (Jacquot et al., 2010; Schroer & Verma, 2021).

Interacting with subpopulations of dynactin that anchor MTs at organelles could allow Dre1 to recruit these organelles to the inclusion, thereby modulating their function. Our work demonstrates that Dre1 specifically localizes to the centrosomal MTOC (Figure 2) and that Dre1 is necessary for *C. trachomatis* to specifically disrupt positioning of MTOC-associated structures, including the centrosome, mitotic spindle, primary cilium, and GA (Figures 3-5). We show that Dre1 contributes to GA recruitment to the inclusion, although it is not exclusively required. Indeed, several other *C. trachomatis* effectors, including InaC/CT813 and ChlaDub1, have been shown to mediate GA fragmentation and recruitment to the inclusion (Kokes et al., 2015; Pruneda et al., 2018; Wesolowski et al., 2017). Thus, multiple host organelles with dynactin-mediated MT organizing capacity are repositioned during infection by Dre1. Furthermore, Dre1 appears dispensable for positioning organelles that lack stable pools of dynactin, such as mitochondria (Figure S5).

Our data demonstrate that Dre1 recruits centrosomes away from the nucleus to the inclusion and prevents clustering of supernumerary centrosomes (Figure 3). Interestingly, the degree to which the centrosome and inclusion associate during infection varies amongst different species of *Chlamydia* and correlates with conservation of *C. trachomatis* Dre1 (Brown et al., 2014; Mital & Hackstadt, 2011; Stephens et al., 1998). Our observation that Dre1 is not required for centrosome overduplication (Figure S3) is consistent with published work that links this process to the *Chlamydia* secreted protease, CPAF (Brown et al., 2014; K. A. Johnson et al., 2009; Knowlton et al., 2011). Our results suggest that *C. trachomatis* Dre1 overrides the host’s centrosome positioning pathways (Figure 3), which results in construction of abnormal spindles (Figures 3, 4). Abnormal spindles interfere with cytokinesis and induce multinucleation, which is a hallmark of *C. trachomatis* infected cells. We demonstrate that loss of Dre1 decreases levels of infection-induced multinucleation (Figure 4), although it does not completely return multinucleation to levels seen in uninfected cells (Figure S4), a finding consistent with work showing that CPAF, and potentially other effectors, also contribute to multinucleation during infection (Alzhanov et al., 2009; Brown et al., 2014). As multinucleated cells have an increased GA content (Sun et al., 2016), we speculate that blocking cytokinesis might be a mechanism by which *C. trachomatis* avoids giving up resources and “real estate” to a host’s daughter cell.

The ability of Dre1 to induce centrosome and mitotic abnormalities could be the key to explaining the role of *C. trachomatis* as a co-factor, along with human papilloma virus (HPV), in the development of cervical cancer (Smith et al., 2004; Wang et al., 2021; Zhu et al., 2016), the fourth leading cause of cancer deaths among women (Arbyn et al., 2020; Jemal et al., 2011). Recent work has identified that HPV and *C. trachomatis* infection induce centrosome overduplication in an additive manner (Wang et al., 2021). As centrosome de-clustering agents have been identified as promising targeted cancer therapeutics (Liu & Pelletier, 2019), Dre1 may be an attractive candidate for future studies. We note that as our experiments thus far have been performed in transformed cell lines, it will be important to test the downstream consequences of Dre-mediated centrosome and mitotic abnormalities in primary cells and in models of HPV infection.

Our work adds to the growing body of evidence that centrosome association is essential for *C. trachomatis* infection. Many Incs, including CT101, CT222, IPAM, CT224, CT228, IncB, IncC, CT288, and CT850 localize at discrete microdomains on the inclusion membrane that are enriched in cholesterol, active Src-family kinases and MTs (Mital et al., 2010; Weber et al., 2015), and co-localize with the centrosome (Mital et al., 2010). Of the Incs found at these microdomains, many have been shown to directly or indirectly bind centrosomal proteins (Almeida et al., 2018; Andersen et al., 2021; Dumoux et al., 2015). Given the reduced genome size of *C. trachomatis*, this represents a striking number of effectors associated with a single organelle. Although we do not know whether endogenous Dre1 is found within inclusion microdomains given the lack of reliable antibodies, our results demonstrate that Dre1 is required to reposition centrosomes around the inclusion membrane during infection.

A unique aspect of *C. trachomatis* infection is that in cells infected with multiple bacteria, individual inclusions traffic to the MTOC and undergo homotypic fusion. These fusion events are critical for pathogenicity (Geisler et al., 2001). IncA, which possesses two SNARE-like domains, is absolutely required for inclusion fusion, but the role of other effectors and host cell components is incompletely understood (Cingolani et al., 2019; Weber et al., 2016). Our data demonstrate that Dre1 contributes to efficient fusion of inclusions and that dynactin may be localized to the site of membrane fusion (Figure 6). We hypothesize that this is another example of Dre1 binding dynactin to position organelles – this time the organelle in question is the inclusion itself. Dre1 does not affect IncA expression or localization, nor does it affect inclusion trafficking to the peri-nuclear region (Figure 1). Rather, our work suggests that Dre1, by targeting centrosomal dynactin, is critical to juxtaposing inclusions, allowing the *C. trachomatis*-encoded fusion machinery to engage. This idea is further supported by our observations that inclusions in L2:L2Δ*dre1* co-infected cells fail to fuse if they are localized on opposite sides of the nucleus (Figure 6). Without Dre1, inclusions may not efficiently discriminate between un-anchored MT minus near the nucleus and the centrosomal MTOC of the host cell. Our results are consistent with data showing that inclusion fusion is delayed when MT minus ends are un-clustered (Richards et al., 2013). Our data add to the literature suggesting a stepwise pathway of establishing fusion-competent inclusions.

We note that we performed the majority of our experiments in HeLa cells, where the MTOC forms at the centrosome, however, *C. trachomatis* typically infects polarized epithelial cells where there are multiple non-centrosomal MTOCs during interphase. Determining how Dre1 perturbs cells where the function of centrosome and the MTOC are separated may also provide insight into the relationship between various classes of MTOCs.

An important question for the future is how Dre1 interacts with dynactin at a molecular level. One model is that Dre1 functions as a mimic for one or more adaptor proteins associated with the dynactin-dynein complex. Most identified adaptors that specify cargo-binding or modulate processivity of the dynactin-dynein complex contain long coiled-coil domains (Reck-Peterson et al., 2018). While we have not yet addressed this question, we note that bioinformatic analysis of Dre1 failed to reveal any sequence or structural homology to known proteins, including known adaptors. Here we demonstrate the dynactin binding domain of Dre1 specifically targets ectopically expressed Dre1 to the centrosomal MTOC and not to other subcellular pools of dynactin (Figure 2). Thus, rather than functioning as a cargo mimic, Dre1 may target specific subpopulations of dynactin that are found at sites where MT minus ends converge. Future structural analyses to determine if Dre1 can distinguish and selectively interact with dynactin in complex with particular adaptors or structures, and to discover whether association with Dre1 alters the regulatory state or activity of dynactin will be required to address these important biological questions.

In summary, we have identified a *C. trachomatis* effector that binds host dynactin, not to facilitate intracellular transport of the pathogen, but rather to reposition organelles including the centrosome and GA around the growing inclusion. Our results suggest a mechanism whereby Dre1 specifically targets dynactin subpopulations that function to cross-link MTs to various organelles and cellular structures. This strategy would allow *C. trachomatis* to override host mechanisms for organelle positioning and create a replicative niche without globally altering organelle function. Our work further highlights how a single pathogen effector can facilitate large scale changes in host cell architecture by interacting with a single, ubiquitous host protein complex. Future elucidation of the molecular basis of Dre1:dynactin interaction may provide insight into the regulation and activities of this essential host protein complex and will have broad implications throughout biology.

## Author Contributions

J.S., C.E., and J.E. conceived, designed, and analyzed experiments. Manuscript was written by J.S. with input from J.E., C.E., R.V., D.S., and N.K. D.S. and J.J. performed MS and bioinformatic analyses. J.S., L.D., E.M., C.E., and R.B. conducted experiments. R.B. and R.V. provided reagents and advice.

## Acknowledgements

We thank Drs. Deborah Dean, Ted Hackstadt, Mary Weber, Dan Rockey, Isabel Derre, Sophie Dumont, and Ron Vale for reagents; Dr. Anita Sil for the use of her confocal microscope; members of the Engel, and Dumont labs for advice, and Drs. Sophie Dumont and Rick McKenney for consulting on the project at various stages. We acknowledge grant support from the NIH (J.S. F31AI133951, J.E. AI073770 and AI105561, L.D. F32AI138372, D.L.S. AI152526, R.V. AI134891, N.J.K. AI122747) and the NSF (J.S. GRFP Fellowship).

## SUPPLEMENTAL FIGURES

**Figure S1.**
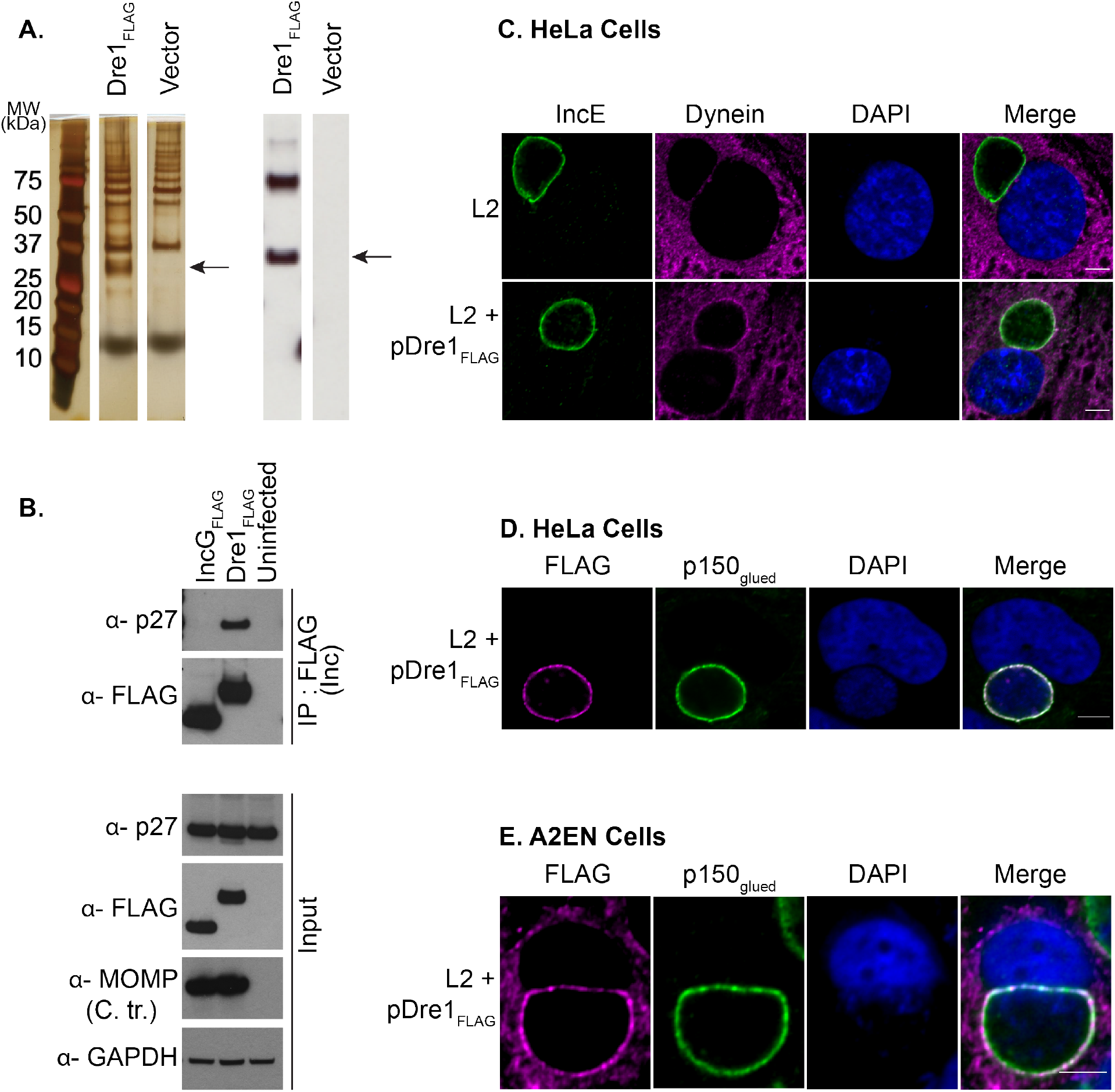
Dre1 interacts specifically with dynactin during infection. (A) Silver stain and anti-FLAG immunoblot of HeLa cells infected with L2+pDre1_FLAG_or L2+vector for 36 hours. Arrows indicate Dre1_FLAG_ monomer bands. The higher bands in the anti-FLAG immunoblot may represent multimeric forms of Dre1_FLAG_. (B) FLAG immunoprecipitations of uninfected HeLa cells, or HeLa cells infected for 36 hours with L2 strains transformed with pDre1_FLAG_, pIncG_FLAG_. Eluates and total lysates were immunoblotted with the indicated antibodies. (C) HeLa cells were infected with L2+pDre1_FLAG_ or L2 for 24 hours, fixed, stained with antibodies specific to dynein (DIC 74.1kDa, magenta) and IncE (green), and counter-stained with DAPI. (D) HeLa cells or (E) pseudo-polarized A2EN cells were infected with L2+pDre1_FLAG_ for 24 hours, fixed, stained with antibodies specific to endogenous p150^glued^ (a dynactin subunit, green) and FLAG (to visualize Dre1, magenta), and counter-stained with DAPI. The anti-FLAG antibody exhibits considerably higher background staining in A2EN cells compared to HeLa cells. Shown are single Z slices. Scale bar, 5µm.

**Figure S2.**
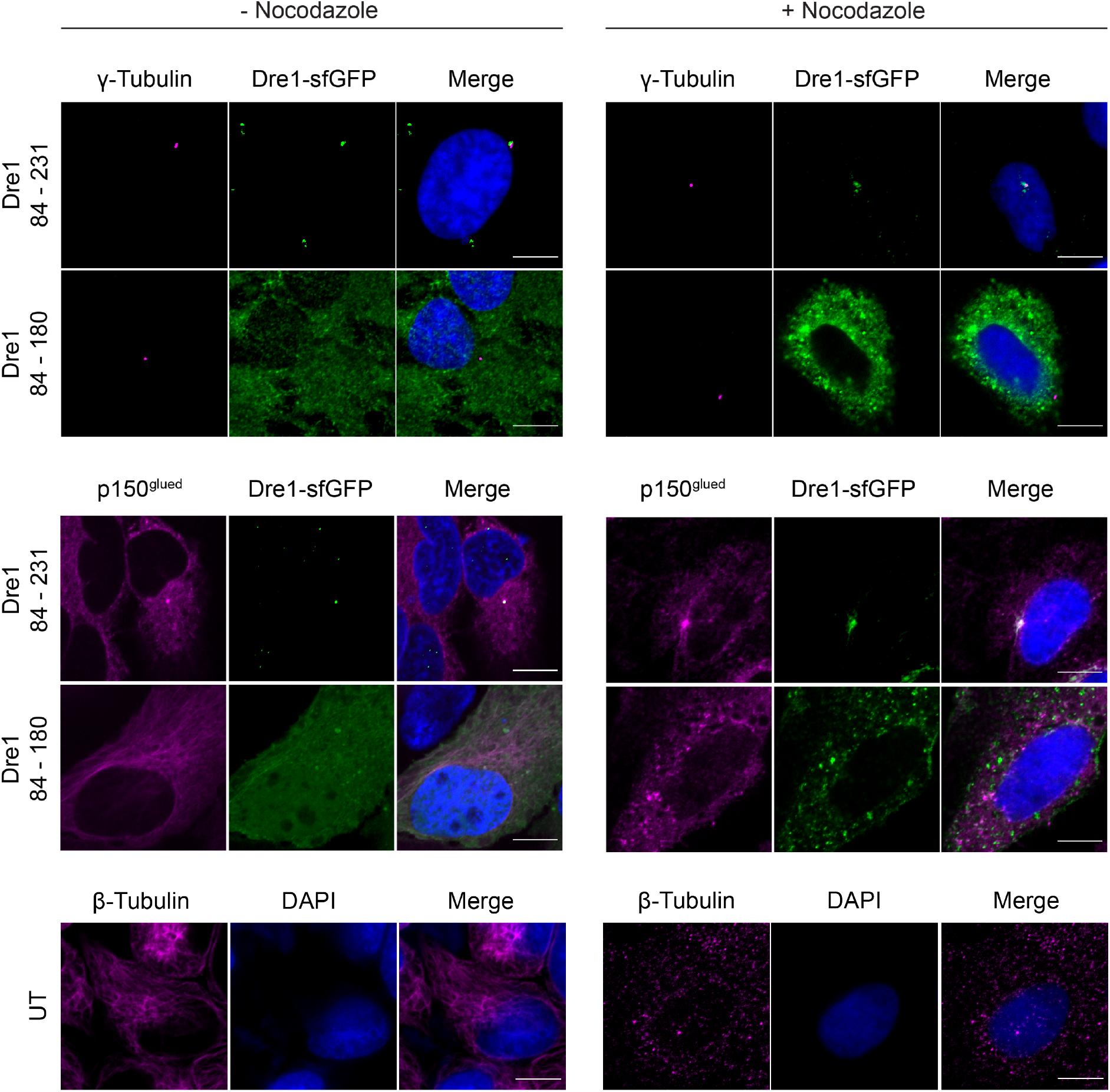
Dre1 localization at the centrosome does not depend on an intact MT network. HeLa cells were transfected for 24 hours with dynactin-binding (amino acids 84-231) or non-binding (amino acids 84-180) variants of Dre1_sfGFP_, exposed to nocodazole (100 ng/mL) or DMSO for three hours, cold-shocked for 30 min on ice, and immediately fixed with 4% PFA. Cells were stained with antibodies to either γ-Tubulin, or p150^glued^ (magenta), and counter-stained with DAPI. To confirm MT depolymerization, untransfected (UT) HeLa cells were treated with nocodazole, cold-shocked, fixed, stained for β-tubulin (to visualize MTs, magenta) and counter-stained with DAPI. Shown are single Z slices. Scale bar, 10µm.

**Figure S3.**
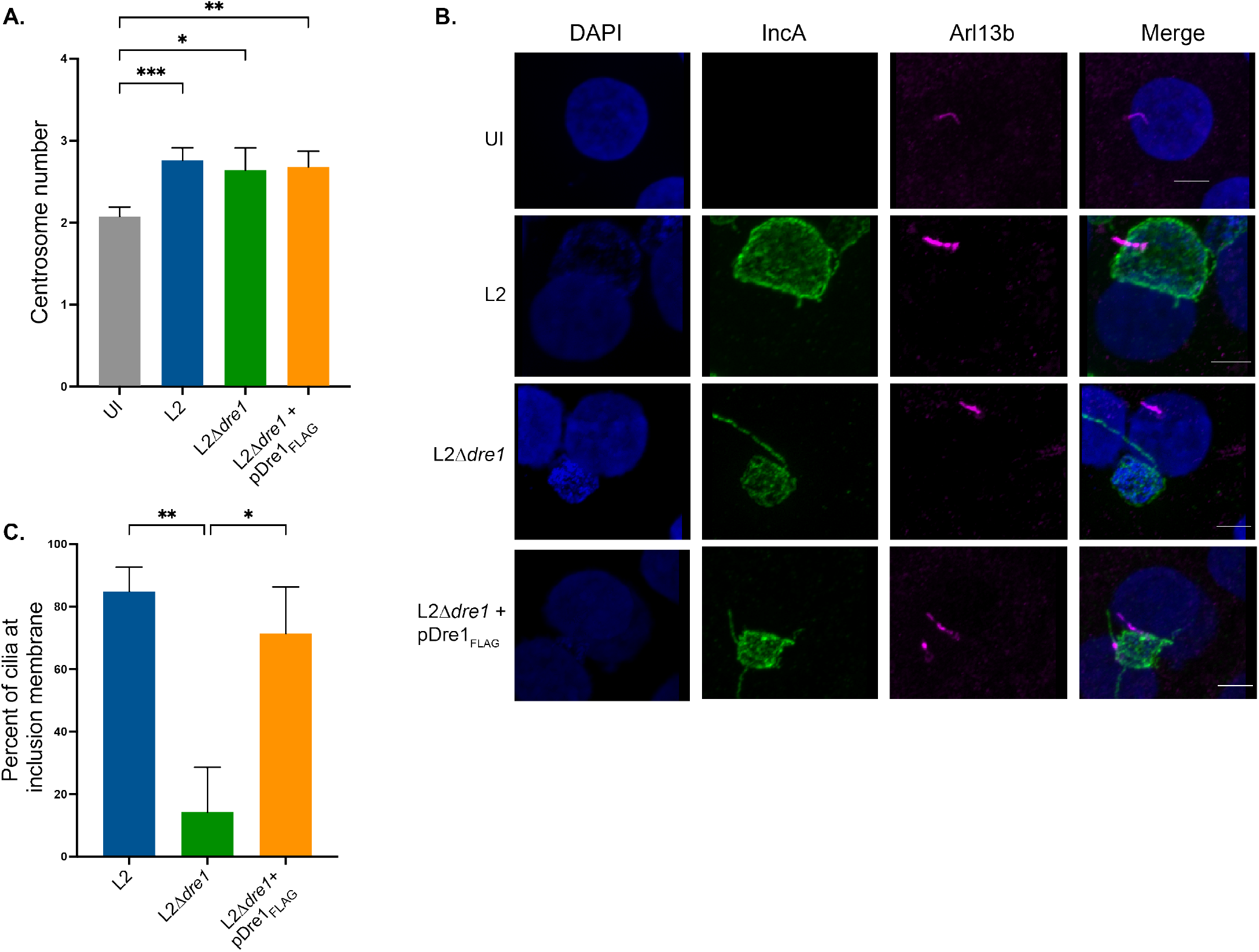
Dre1 modulates the position of primary cilia during infection but is not involved in *C. trachomatis*-induced dysregulation of centrosome duplication. (A) HeLa cells were infected with the indicated L2 strains for 36 hours, fixed, stained with antibodies to γ-Tubulin and Centrin to visualize centrosomes, MOMP to visualize bacteria, and counter-stained with DAPI. The average number of centrosomes in cells ± SD (n=3) is reported. 100 cells per condition per replicate were scored. (B) Pseudo-polarized A2EN cells were infected with the indicated L2 strains for 24 hours, fixed, stained with antibodies to Arl13b to visualize primary cilia (magenta) and IncA to visualize the inclusion membrane, and counter-stained with DAPI. Shown are single Z slices. Scale bar, 10µm. In some images, previously described IncA-positive tubules extending from the inclusion are present (Mirrashidi et. al., 2015) and are clearly distinct from the Arl13b-positive primary cilium (C) The percentage of cilia anchored at the inclusion membrane from the four different conditions is shown. 25 ciliated cells per condition were analyzed, over two biological replicates. Data are represented as mean ± SD. *p<0.05, **p<0.01, ***p<0.001, Welch’s ANOVA.

**Figure S4.**
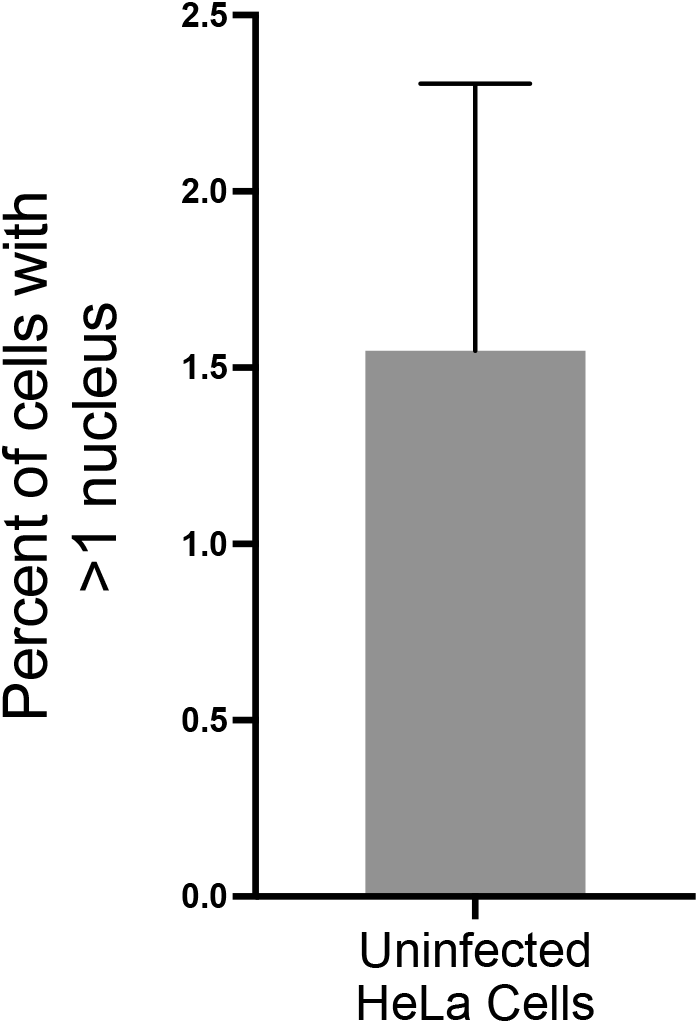
Basal rate of multinucleation in uninfected HeLa cells. HeLa cells were seeded on coverslips and 48 hours later fixed and stained with DAPI to visualize nuclei and with WGA 647 to delineate the plasma membrane. Approximately 1.5% of uninfected HeLa cells contained more than one nucleus. 120 cells were analyzed, over three biological replicates. Data are represented as mean ± SD.

**Figure S5.**
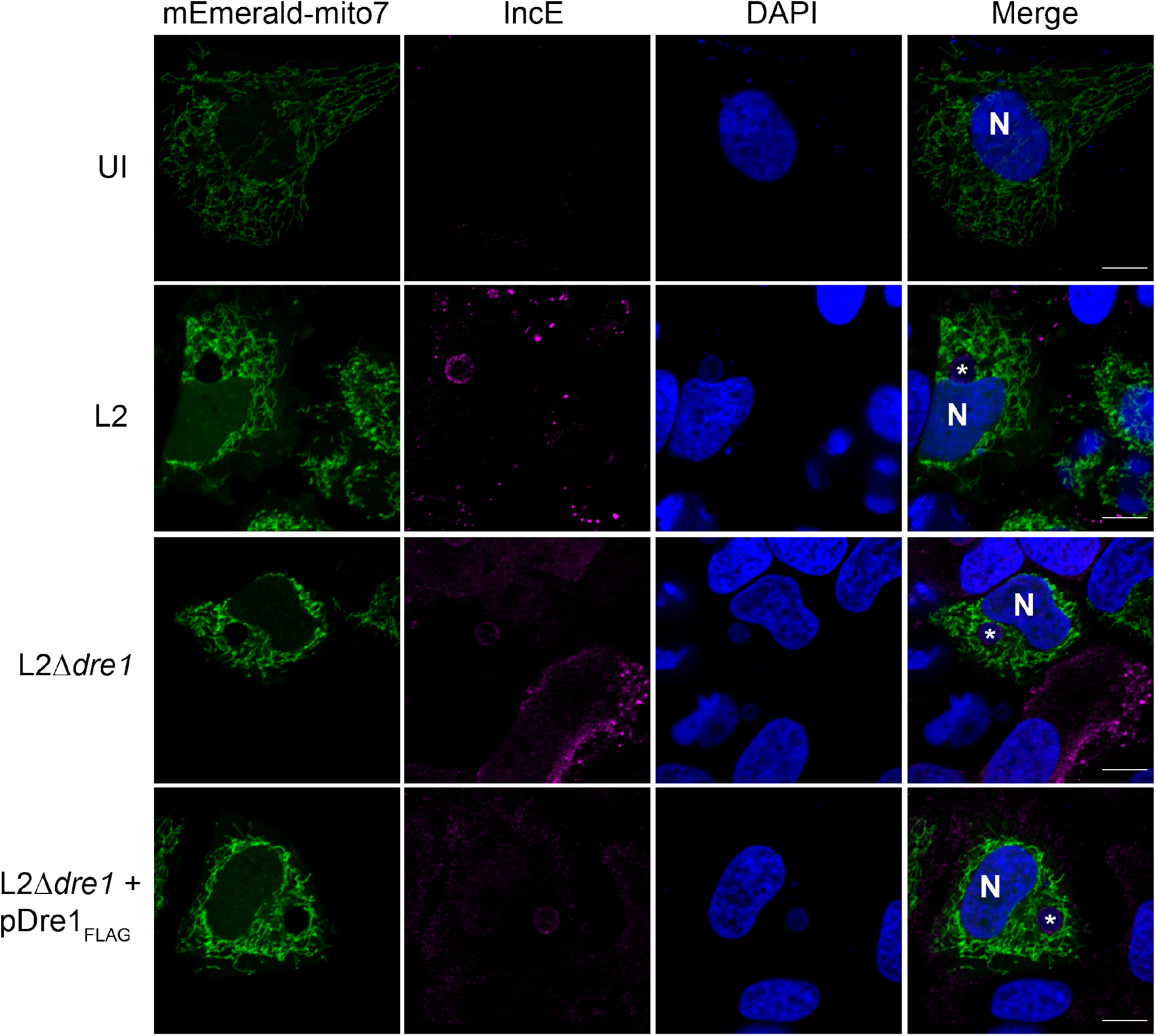
Dre1 does not alter mitochondrial position during infection. HeLa cells were transfected with mEmerald-Mito7 (a mitochondrial marker, green), infected for 24 hours with the indicated L2 strains, fixed, stained with an antibody specific to IncE (to visualize the inclusion membrane, magenta), and counter-stained with DAPI. N, nucleus. *, inclusion. Shown are single Z slices. Scale bar, 10µm.

**Figure S6.**
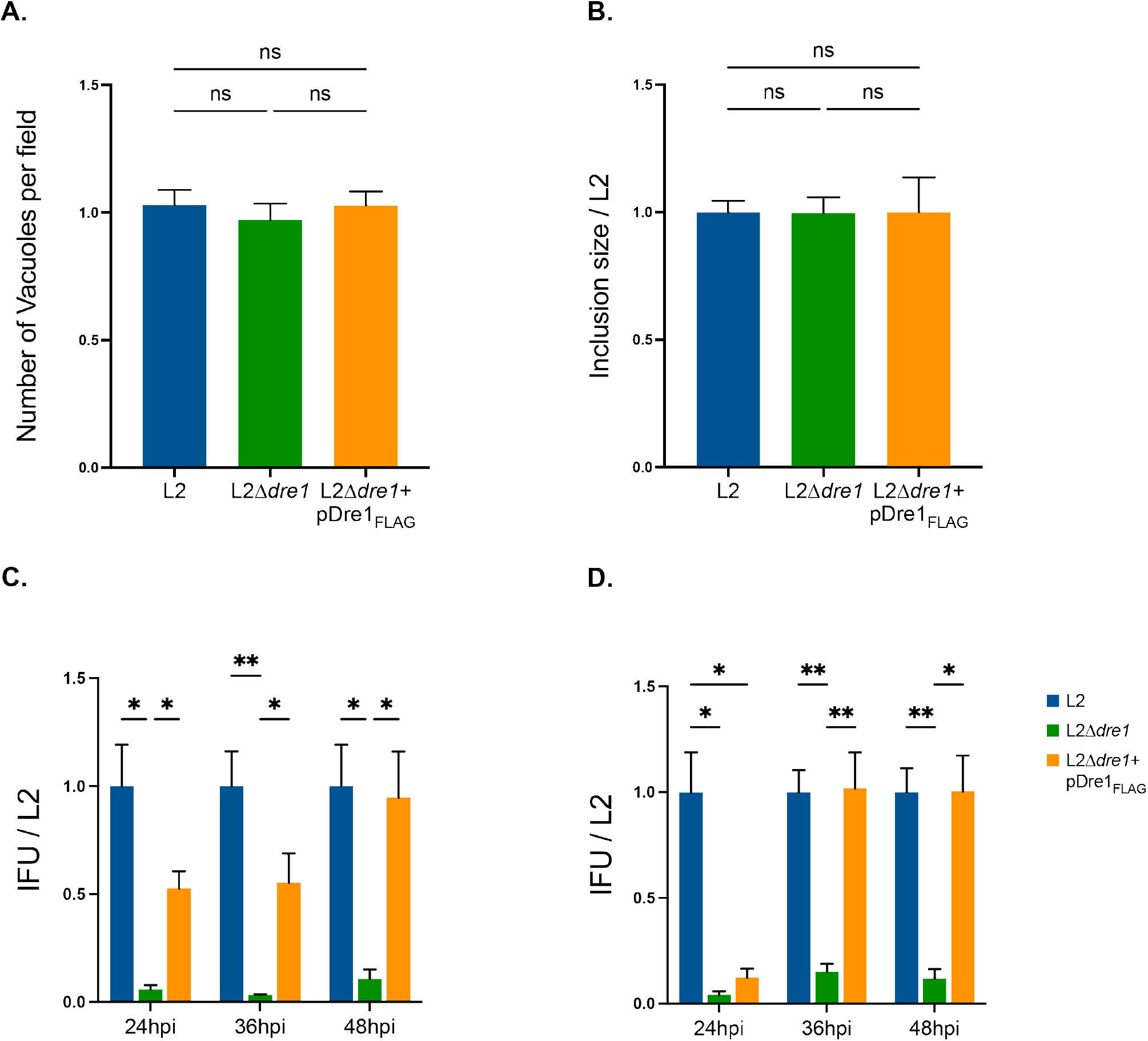
L2Δ*dre1* exhibits a growth defect at later stages of the infectious cycle. (A, B) Quantitation of (A) inclusion number and (B) inclusion area in HeLa cells infected with the indicated L2 strains for 24 hours. For (A), 35 fields per replicate for each condition were counted. For (B) 3D reconstructions for 35 inclusions per replicate for each condition were generated and area of each reconstructed inclusion was measured using Fiji. Inclusion area for each strain was normalized to L2 inclusion area. Data are mean ± SD of three independent experiments. (C, D) Quantitation of infectious progeny at 24, 36, and 48 hpi in (C) pseudo-polarized A2EN cells or in (D) HeLa cells infected with the indicated L2 strains. Relative IFUs for 24, 36, and 48 hpi are expressed as a fraction of L2 infection at each time point. Data are mean ± SD from 4 independent experiments. *p<0.05, **p<0.01, ***p<0.001, Welch’s ANOVA.

## Tables

Supplementary Table 1

**Scoring for all PPIs identified through AP-MS (Transfection and Infection Interactomes).**

Related to Figure 1 and S1.

Worksheet 1, Transfection Interactome (all). Summary of MiST and CompPASS scores from Dre1_Strep_-prey PPIs detected by AP-MS as well as individual MiST scores for abundance, reproducibility, and specificity. From Mirrashidi et. al., 2015. Worksheet 2, Transfection Interactome (High Confidence; HC). Same as Worksheet 1, except that it only contains data for the high-confidence Dre1-host PPIs (as defined in Mirrashidi et. al., 2015.) Worksheet 3, Infection Interactome (all). Spectral counts and SAINT scores with associated Bayesian False Discovery Rate scores (BFDR) for all Dre1_FLAG_-host interactions identified during *C. trachomatis* infection.

Supplementary Table 2

Whole genome sequencing identifying SNVs in *C. trachomatis* L2Δ*dre1* and parental strain compared to the published *C. trachomatis* L2 434/Bu genome sequence (accession no. AM884176).

Supplementary Table 3

List of primers and strains used in this study.

